# On-scalp MEG SQUIDs are sensitive to early somatosensory activity unseen by conventional MEG

**DOI:** 10.1101/686329

**Authors:** Lau M. Andersen, Christoph Pfeiffer, Silvia Ruffieux, Bushra Riaz, Dag Winkler, Justin F. Schneiderman, Daniel Lundqvist

**Affiliations:** NatMEG, Department of Clinical Neuroscience, Karolinska Institutet, Nobels väg 9, 171 77, Stockholm, Sweden; Department of Microtechnology and Nanoscience – MC2, Chalmers University of Technology, SE-412 96, Gothenburg, Sweden; MedTech West and the Institute of Neuroscience and Physiology, Sahlgrenska Academy, University of Gothenburg, Gothenburg, Sweden

## Abstract

Magnetoencephalography (MEG) has a unique capacity to resolve the spatio-temporal development of brain activity from non-invasive measurements. Conventional MEG, however, relies on sensors that sample from a distance (20-40 mm) to the head due to thermal insulation requirements (the MEG sensors function at 4 K in a helmet). A gain in signal strength and spatial resolution may be achieved if sensors are moved closer to the head. Here, we report a study comparing measurements from a seven-channel on-scalp SQUID MEG system to those from a conventional (in-helmet) SQUID MEG system.

We compared spatio-temporal resolution between on-scalp and conventional MEG by comparing the discrimination accuracy for neural activity patterns resulting from stimulating five different phalanges of the right hand. Because of proximity and sensor density differences between on-scalp and conventional MEG, we hypothesized that on-scalp MEG would allow for a more high-resolved assessment of these activity patterns, and therefore also a better classification performance in discriminating between neural activations from the different phalanges.

We observed that on-scalp MEG provided better classification performance during an early post-stimulus period (15-30 ms). This corresponded to electroencephalographic (EEG) response components N16 and P23, and was an unexpected observation as these components are usually not observed in conventional MEG. They indicate that on-scalp MEG opens up for a richer registration of the cortical signal, allowing for sensitivity to what are potentially sources in the thalamo-cortical radiation and to quasi-radial sources.

We had originally expected that on-scalp MEG would provide better classification accuracy based on activity in proximity to the P60m component compared to conventional MEG. This component indeed allowed for the best classification performance for both MEG systems (60-75%, chance 50%). However, we did not find that on-scalp MEG allowed for better classification than conventional MEG at this latency. We believe this may be due to the limited sensor coverage in the recording, in combination with our strategy for positioning the on-scalp MEG sensors. We discuss how sensor density and coverage as well as between-phalange source field dissimilarities may influence our hypothesis testing, which we believe to be useful for future benchmarking measurements.

## Introduction

On-scalp magnetoencephalography (MEG) holds great promise for improving MEG in terms of better spatial resolution. In this study, we investigated this promise by comparing the spatial resolution of on-scalp MEG recordings to that of conventional state-of-the-art MEG.

On-scalp MEG technology is becoming increasingly available through the use of high critical temperature (operating at ∼77 K) superconducting quantum interference devices (high-*T*_c_ SQUIDs) (Faley et al., 2013; Öisjöen et al., 2012; Zhang et al., 1993) and optically pumped magnetometers (OPMs) (Boto et al., 2017; Budker and Romalis, 2007). The present on-scalp sensor technology, though, has significantly higher noise levels than the state-of-the-art MEG sensors, which operate at a low critical temperature (low-*T*_c_). Hence, while low-*T*_c_ SQUIDs (hereafter called in-helmet sensors) demonstrate noise levels of ∼3 fT/vHz, the very best high-*T*_c_ sensors show noise levels of ∼10 fT/vHz (Faley et al., 2006) and often near 50 fT/vHz (Pfeiffer et al., 2019b). These differences in noise levels mean that a trade-off might be expected between on-scalp and in-helmet sensors, with on-scalp sensors promising the highest gain in signal-to-noise ratio (SNR) for superficial sources (Iivanainen et al., 2017; Riaz et al., 2017).

A fundamental component of neuroscience research is to elucidate the functional organization of the brain by resolving and discriminating patters of brain activity. EEG has historically played - and plays - a pivotal part in revealing especially the temporal properties of these activations. The capacity to spatio-temporally resolve such responses, however, improved markedly with the advent of MEG (Hari et al., 1984; Tiihonen et al., 1989), offering a temporal resolution of <1 ms, and spatial resolution on the millimetre level. Note that the spatial resolution, however, depends on a multitude of factors, such as the nature and location of the sources underlying an activity pattern, the distance between the sources and the sensors, the source strength, etc. (Hari and Puce, 2017). Conventional state-of-the-art MEG sensor arrays are typically housed inside a fixed-size helmet and located 20-40 mm away from the scalp of subjects due to the thermal insulation needed to keep the sensors at operational temperature (∼4 K). Consequently, because the magnetic field magnitude is roughly inversely proportional to the squared distance between a source and a sensor, on-scalp MEG arrays, wherein magnetometers can be positioned within a few millimetres of the scalp surface, hold great promise for registering weak, but shallow, neural sources (Iivanainen et al., 2017).

A specific type of shallow sources that on-scalp MEG may be sensitive to is “radial” sources in the cortex. This seems to be in conflict with the fact that the radial component of a magnetic field detected by MEG sensors on the outside of a conducting *sphere* is zero for sources that are radial. The head is only *quasi*-*spherical*, however (Hämäläinen et al., 1993). Near the primary somatosensory cortex (S1), the head is in fact not well approximated by a sphere. The less spherical the head is, the less meaningful it thus is to speak of radial sources. In these cases, it would be more meaningful to speak of quasi-radial sources, which may produce a non-zero magnetic field outside the head. It is thus probable that on-scalp MEG may be sensitive to quasi-radial sources that are on or near the crests of the brain’s gyri because of the close proximity between them and the sensors (∼1 cm), as compared to conventional MEG (∼3+ cm).

Another type of source that on-scalp MEG may be sensitive to are sources from some of the white matter fibres. Specifically, the thalamo-cortical radiation (between thalamus and S1) is a potential candidate. Currents within white matter axons can be modelled by quadropoles, which can be seen as two dipoles with opposite polarities. This makes them invisible when recorded from a distance. However, where white matter tracts have a bend the two dipoles will not have exact opposite polarities. Signals likely to have arisen from the thalamo-cortical radiation have in fact been measured using EEG at latencies of 15-20 ms post-stimulus (Buchner et al., 1995; Gobbelé et al., 1998), and it has also more recently been localized using a combination of MEG and EEG to the thalamo-cortical radiation (Götz et al., 2014).

Moving onto the scalp also changes the requirements and possibilities for the size of the pick-up coils of magnetometers. Specifically, in order to gain in spatial resolution with on-scalp MEG, high spatial sampling of the neuromagnetic field may be required. The most practical way to achieve this is to reduce the size of the on-scalp pick-up coils, despite a trade-off in terms of increased sensor noise when making pick-up coils smaller. The smaller pickup coils also means that sensors can be packed more closely together than sensors with bigger pickup coils, such as the in-helmet sensors. The combination of sensor proximity and smaller sensor size means that it should be possible to spatially separate neural activity more finely with on-scalp MEG than in-helmet MEG.

To design a test for ascertaining whether on-scalp MEG provides a finer spatial resolution than in-helmet MEG, we devised a stimulation sequence where five different phalanges with varied between-phalange proximity were stimulated on the same hand. The distal phalange of the little finger, the index finger, and the thumb were chosen alongside the middle and proximal phalanges of the index finger. It has been shown that MEG signals can be separately identified from different finger representations in the sensory homunculus (Baumgartner et al., 1991). There is also evidence of the phalanges being separately represented in the sensory homunculus (Druschky et al., 2002).

Our rationale was thus that, due to differences in somatotopic representation, the three phalanges on the index finger and the thumb would be the hardest to discriminate from one another whereas the little finger should more easily be separated from the rest (Baumgartner et al., 1991; Druschky et al., 2002). Because of the gain in sensitivity to magnetic fields, and because of the higher density of sensors, we expected that on-scalp MEG would have an advantage over conventional MEG in terms of discriminating between the activity patterns of these superficial sources that are all close to one another. Tactile stimulations are known to result in a volley of responses with early (< 25 ms) and late (> 50 ms) components, each of which has a spatio-temporal trajectory evolving along time-courses on the order of milliseconds, as has been revealed by electroencephalography (EEG) (Tsuji and Murai, 1986; Yamada et al., 1984). We expected the classification to be maximal around the P60m since this is the strongest of the primary somatosensory responses. Note that this was not a test devised for detecting quasi-radial sources or the thalamo-cortical radiation.

### Snapshot of results

P60m did indeed allow for the best classification performance for both MEG systems (60-75%, chance 50%). However, we did not find that on-scalp MEG allowed for better classification than conventional MEG. We provide more details on this in the methods and results sections below. We also observed that on-scalp MEG provided better classification performance during an early post-stimulus period (15-30 ms). This corresponded to electroencephalographic (EEG) response components N16 and P23. These were unexpected observations as P23 is considered to be generated by radially oriented sources, which theoretically should be invisible to MEG as discussed above. Furthermore, N16 is most likely related to the thalamo-cortical radiation from thalamus to S1.

## Methods

### Subjects

Four subjects were recruited from the scientific team involved in the recordings (four males; aged 49 y, 39 y, 34y and 30 y). The experiment was approved by the Swedish Ethical Review Authority (DNR: 2018/571-31/1), in agreement with the Declaration of Helsinki.

### Stimuli and procedure

The tactile stimulations were generated by using five inflatable membranes (MEG International Services Ltd., Coquitlam, Canada) fastened to the subject’s right hand. The membranes were placed on the distal phalange of the little finger (L1), the distal phalange of the thumb (T1) and the three phalanges of the index finger (distal phalange: I1, middle phalange: I2, proximal phalange: I3) (Fig. 1A). The membranes setup was part of a custom stimulation rig (built by Veikko Jousmäki, Aalto University, Finland), which was controlled by pneumatic valves (model SYJ712M-SMU-01F-Q, SMC Corporation, Tokyo, Japan) using 1 bar of pressurised air.

**Fig. 1:**
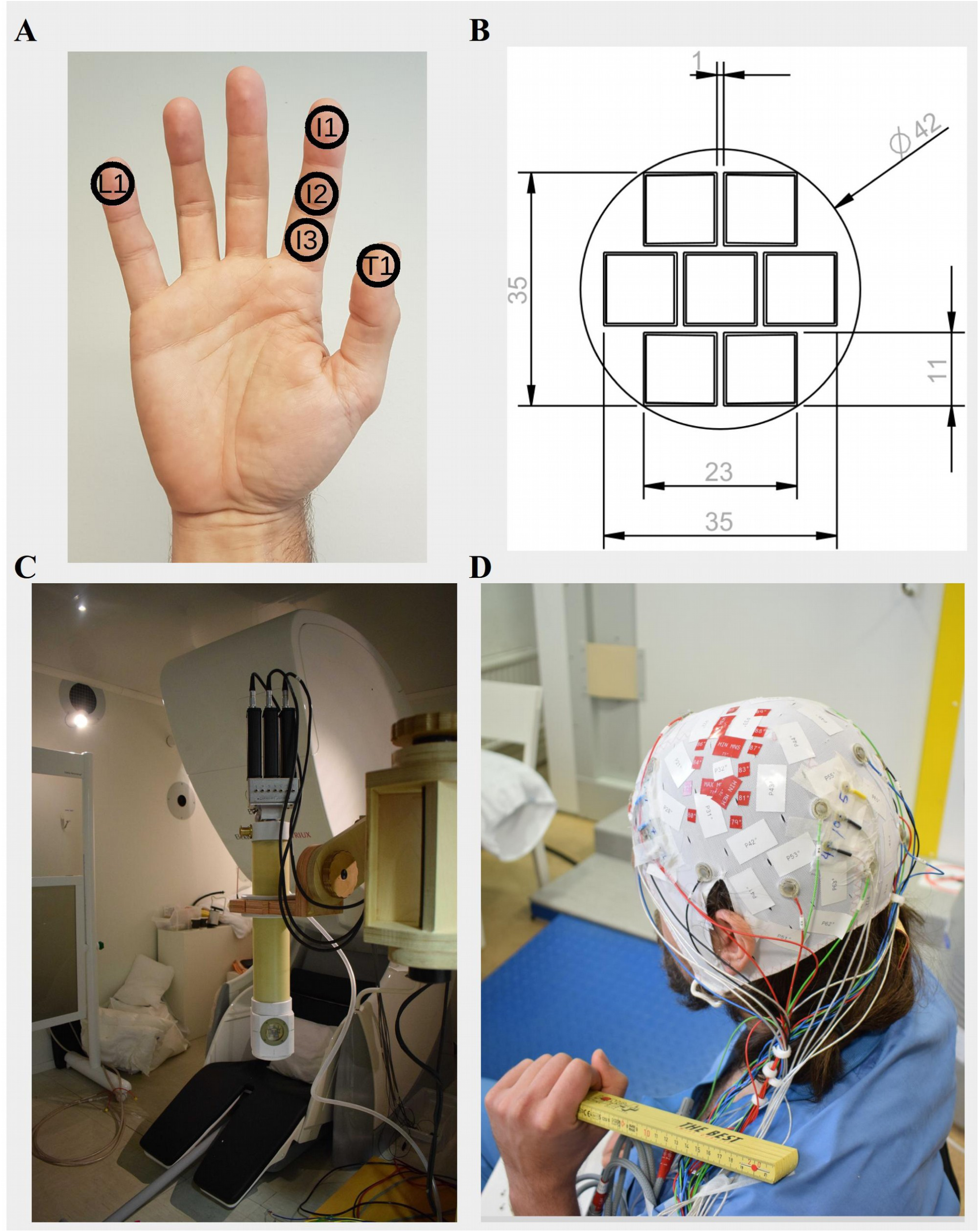
Illustrations of equipment used. **A:** The five phalanges stimulated **B:** Schematic layout of the sensor array. All measurements are in millimetres. **C:** The on-scalp MEG system cryostat on a wooden articulated armature in front of the in-helmet MEG system. **D:** The customised cap prepared on a subject. The white vinyl markers correspond to the seventy-four channels on the EasyCap 74 channel layout. The larger red vinyl markers indicate the maximum and minimum positions to which the on-scalp MEG was aimed. The smaller red vinyl markers show the fifteen additional coordinate points used to assist and optimize the positioning. Nineteen EEG electrodes were attached in a 10-20 montage (positions Fp1, Fp2, F7, F3, Fz, F4, F8, T7, C3, Cz, C4, T8, P7, P3, Pz, P4, P8, O1 and O2). EEG electrodes that would hinder scalp contact during on-scalp MEG placement were removed at the point of measurement. Nine HPI-coils were placed around the planned recording positions

The experimental paradigm consisted of 5,000 stimulations evenly distributed between the five phalanges (1,000 stimulations each). The inter-stimulus interval (ISI) was 333 ms. Stimulations were ordered in blocks of 10 where the order of stimulations within a block was pseudo-random such that each stimulation type occurred two times in each of these blocks. There was a break in the stimulation sequence every 1,250 trials (four total). At the beginning of the experiment, at each break and at the end of the experiment, the sleepiness level of the subject was assessed using the Karolinska Sleepiness Scale KSS (Åkerstedt and Gillberg, 1990). During the stimulation procedure, subjects were watching a television show of their own choice to keep them at a stable level of wakefulness. Audio was provided via sound tubes (model ADU1c, KAR Oy, Helsinki, Finland) rendering the tactile stimulation inaudible. Subjects were instructed to pay full attention to the show and to pay no attention to the stimulations. The subjects held the stimulated hand under the table such that the hand could not be seen.

The experimental paradigm was run three times on each subject, once using conventional in-helmet MEG (Elekta TRIUX) with 102 magnetometers and 204 planar gradiometers; and two times with a seven channel high-*T*_c_ SQUID-based on-scalp MEG system (hereafter referred to as the “on-scalp MEG”) sampling from two head positions per subject. During all three recordings, 19 EEG electrodes in a customized 10-20 montage were used to record electrical potentials. The in-helmet MEG was done as a conventional whole-head recording, both for comparison, but also for guiding placement of the on-scalp MEG system.

To project the likely positions of the magnetic field maximum and minimum onto the subjects’ heads, an initial localizer protocol was run in in-helmet MEG, using 1,000 stimulations to the upper phalange of the index finger (I1). Equivalent Current Dipoles (ECDs) were then fitted to the P60m component of the responses to these stimulations, for each sample point between 45 ms and 65 ms based on the activity of the 102 magnetometers. The magnetic field pattern generated by the dipole with the least residual variance was subsequently projected onto the scalp of the subject using a single compartment volume conduction model based on the subject’s anatomy. The centres of the projected maxima and minima were chosen for the recording position of the centre sensor of the on-scalp MEG system (i.e., the centre of the cryostat window over the high-*T*_c_ magnetometers). This procedure has been applied successfully before (Andersen et al., 2017; Xie et al., 2016).

### Equipment

All measurements were carried out in a two-layer magnetically shielded room (MSR; model AK3b from Vakuumschmelze GmbH & Co. KG, Hanau, Germany) at the NatMEG facility, Karolinska Institutet, Stockholm, Sweden, http://www.natmeg.se/).

For the in-helmet MEG, an Elekta Neuromag TRIUX (Elekta Oy, Helsinki, Finland) was used. The Elekta Neuromag TRIUX contains 102 sensor chips, each with a magnetometer channel with a pickup loop size of 21 mm × 21 mm and two orthogonal planar gradiometer channels.

The 7-channel on-scalp MEG-system was made at Chalmers University of Technology (Pfeiffer et al., 2019b). It contains seven high-*T*_c_ SQUID magnetometers, each with a pick-up loop size of 9.2 mm × 8.7 mm arranged in a hexagonal array (Fig. 1). The seven sensors were mounted on a sapphire window connected to a liquid nitrogen vessel inside a vacuum cryostat, the tail of which has a 0.4 mm thick polymer vacuum tight window in front of the sensors. The cryostat is placed on the subject’s head with the help of a wooden articulated armature (Fig. 1C). The separation between the scalp and the sensors is about 3 mm or less. The output of the SQUID electronics was recorded using the analogue miscellaneous (MISC) channels of the Elekta TRIUX system, which allowed for sampling the on-scalp data using the same clock as for the in-helmet data and also for synchronizing on-scalp MEG recordings to the EEG data.

19 EEG scalp electrodes were also used, placed in a standard 10-20 layout on a customizable 128 channel cap (Cap and electrodes from EasyCap, BrainVision LLC, USA).

### Preparation of subjects

In preparation for the localizer procedure and the three experimental runs, each subject was fitted with a customized EEG-cap. For the purpose of visualizing a grid of coordinates on an individual’s scalp, a total of 89 positions were marked on the 128 channel cap using vinyl markers, from which the first 74 positions were according to the EasyCap 74 channel layout (BrainVision LLC, USA). 15 extra positions were marked out around the area where we expected the P60m field maxima and minima to occur (see Fig. 1), which served to increase the density of coordinates and aid the final positioning of the on-scalp MEG system. In addition, nine Head Position Indicator (HPI) coils were placed around the expected positions of the maximum and minimum field positions while leaving room for placement of the on-scalp MEG system in between them.

A Polhemus FASTRAK system was used to digitize these eighty-nine positions. In addition, three fiducial points were digitized (the nasion, and the left and right pre-auricular points) as well as the HPI coils. Finally, two- to three-hundred extra head shape points were digitized to aid with co-registration. The reference electrode was placed on the subject’s right cheek and the ground electrode on the backside of the neck. For each subject, some of the electrodes had to be removed to leave room for the tail of the cryostat and the HPI-coils. For Subject 1, these were: C3, Cz, P3 and Pz; for Subject 2: C3, Cz and P3; for Subject 3: C3 and Cz; for Subject 4: C3 and Cz.

### Acquisition of data

All data was acquired inside the MSR and was sampled on the Elekta Neuromag TRIUX at a sampling rate of 5,000 Hz with an online band-pass filter between 0.1 Hz and 1,650 Hz. The nine HPI coils were running continuously at frequencies 1,137-1,537 Hz with steps of 50 Hz between them. After recording Subject 1, this was changed for the on-scalp recordings, though, because the HPI-coil frequencies were clearly visible on the raw data. For the three subsequent subjects, HPI coils were activated only semi-continuously during regular breaks (30 s) in the on-scalp recordings. Subjects were motivated, comfortably seated with their heads semi-rigidly fixed relative to the on-scalp MEG system, and instructed to be still. It was thus our judgement that running the HPI coils semi-continuously would be sufficient for applying the sensor localization method of Pfeiffer et al. (2018), as also evidenced in Pfeiffer et al. (2019a).

### Processing of MEG and EEG data

MNE-Python (Gramfort et al., 2013) was used to read in the raw in-helmet MEG, and on-scalp MEG and EEG data. The raw data was subsequently low-pass filtered at 30 Hz. Bad EEG channels, if any, were then marked as bad and the EEG was re-referenced to a common average reference using only the good channels. Subject 1 had no EEG marked as bad; Subject 2 had Fz marked as bad; Subjects 03 and 04 had T8 marked as bad. Among the on-scalp MEG channels, two channels were marked as bad for each subject. For Subject 1: 006 and 007; for Subject 2: 001 and 007; for Subject 3: 001 and 007; for Subject 4: 005 and 007. The filtered data was then segmented into epochs containing data with a 30 ms pre-stimulus period and a 300 ms post-stimulus period. Data was also time-shifted 41 ms to compensate for the delay between the trigger onset and the actual delivery of the pneumatically driven stimulation.

Subsequently, the segmented data was exported to FieldTrip (Oostenveld et al., 2011) where epochs that revealed large variance within them were marked using visually guided tools. This was done for each on-scalp MEG channel and for each of the in-helmet MEG channels. Finally, event-related fields and potentials (ERFs and ERPs) were calculated from the epochs after having removed those with high variance.

For the in-helmet channels, the peak time was identified for each of the individual phalange P60m components. The two magnetometers with the largest positive and negative response at this time point, respectively, were identified as *the extrema magnetometers*. The six magnetometers geodesically closest to each of the two extrema magnetometers were also identified. These two times seven magnetometers were later to be used in the classification comparisons.

### MR-preprocessing

A full segmentation of earlier acquired T1 MR-images was performed with FreeSurfer (Dale et al., 1999; Fischl et al., 1999). Based on this segmentation the watershed algorithm of FreeSurfer was used to extract surfaces indicating the boundaries for the skin, the skull and the brain. A single compartment volume conductor was subsequently created based on the boundary for the brain surface. For each subject, the T1 was co-registered to the subject’s head shape. First, a rough alignment was done using the fiducials. This was subsequently optimized using an Iterative Closest Points algorithm.

### Dipole fits

Finally, a dipole fit was done for each of the five phalanges based on the in-helmet MEG data. A single dipole was fit based on the data from twenty-six magnetometers at and around the extrema, using non-linear optimization for the topographies between 50.0 ms and 70.0 ms in steps of 0.2 ms.

### Classification of phalanges

For classifications of the phalanges, we used logistic regression (Bishop, 2006). Each phalange was compared to every other phalange at each time sample. The aim of the classification was to investigate whether the brain’s responses would be sufficient to discern between which phalange was stimulated.

For the classification, the 200 trials with the highest variance were removed from the 5,000 trials in total. Afterwards, for each of the ten possible comparisons, the number of trials for each phalange was equalized. The data was normalized by subtracting the mean and dividing by the variance before being fed to the logistic regression. It was regularized using the L2 norm. The classifier was trained using cross-validation (ten splits). The Python module *Scikit-learn* was used to implement the classification (Pedregosa et al., 2011).

Optimally, all seven on-scalp magnetometers would have been used for the classifications, but for each subject, the on-scalp recording contained two high-noise magnetometers. For the on-scalp data, these comparisons were hence run using only the data of all five low-noise magnetometers.

To make a fair comparison with the in-helmet data, these comparisons were also run with five in-helmet magnetometers, i.e. the extremum magnetometer and the four geodesically closest neighbours were run. All these analyses were run for the maximum and minimum positions separately.

## Results

In this section, results related to the original hypothesis are presented first, followed by a separate section for the unexpected results related to the early components.

Unless otherwise stated, plots are based on data from Subject 4. This subject is typical in terms of the in-helmet recording and showed some interesting findings in the on-scalp recordings. Plots for all other subjects can be found in the supplementary material.

### Event-related Potentials (ERPs) and Fields (ERFs)

Subjects showed the expected P60m component (Fig. 2, Supplementary Fig. 1). However, within single sensor data, the P60m differed in peak latencies between the phalanges (Fig. 2ADE). It can be seen that the peak topographies are very similar, especially within the three phalanges of the index finger. Amplitudes differ more for the on-scalp compared to the in-helmet recordings though. This is likely to be because the on-scalp sensors sample a smaller area than the in-helmet sensors. For the other two fingers, the dipolar fields are slightly rotated relative to the ones for the index finger (Fig. 2B).

**Fig. 2:**
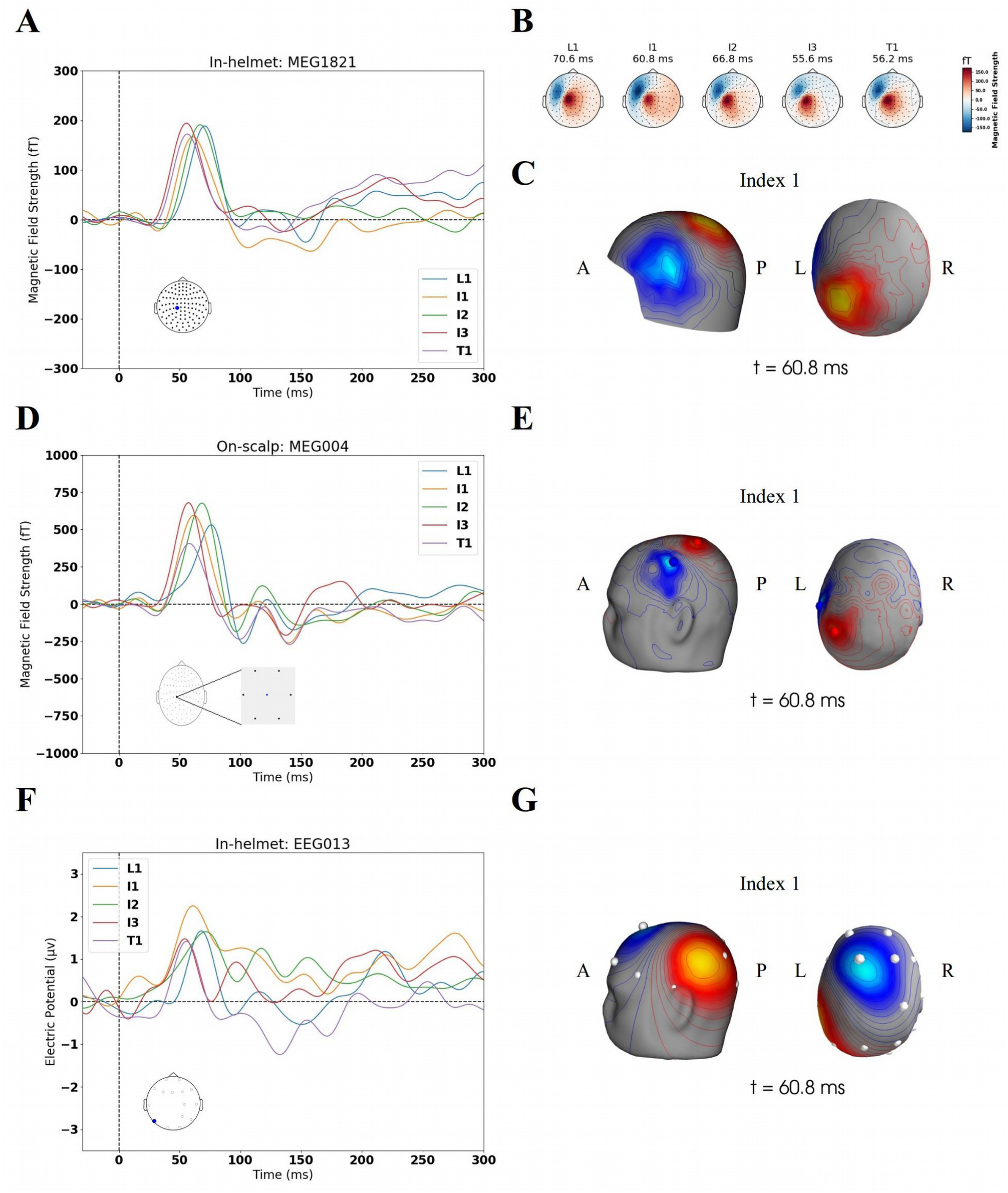
Summary of activation for Subject 4. **A:** Event-related fields on in-helmet magnetometer reflecting the maximum peaks. **B:** 2d-topographies for the time point where MEG1821 (**A**) has its maximum. **C:** 3d-topography for Index 1 (I1) at its peak. **D:** Event-related fields on on-scalp magnetometer placed on the projected maximum. **E:** The fitted dipole’s (Fig. 3) projected magnetic field on the scalp. The blue and red dots are the positions of the centre magnetometer of the cryostat **F:** Event-related potentials on an electrode showing the P60. **G:** The fitted dipole’s (Fig. 3) projected electric potential on the scalp. The white dots are the positions of the electrodes. The P60m (panels A and D) and P60 (panel F) are clearly seen for all phalanges though with different peak times. On the EEG (panel F), the P23 is also seen. See also Supplementary Fig. 1 for all subjects.

### Dipole fits

Using the dipole fitting procedure the P60m components were fitted to the primary somatosensory cortex (S1) with mean goodness of fit over all five phalanges being above 90% for all subjects (Fig. 3A). The greatest differences in dipole positions and dipole orientations were found between L1 and the other phalanges as expected from the somatotopic distance (Fig. 3B). Greater distances between sources and greater differences in orientations also correlated positively with better classification accuracy (Fig. 3C). See Supplementary Figs. 2-6 for all subjects.

**Fig. 3:**
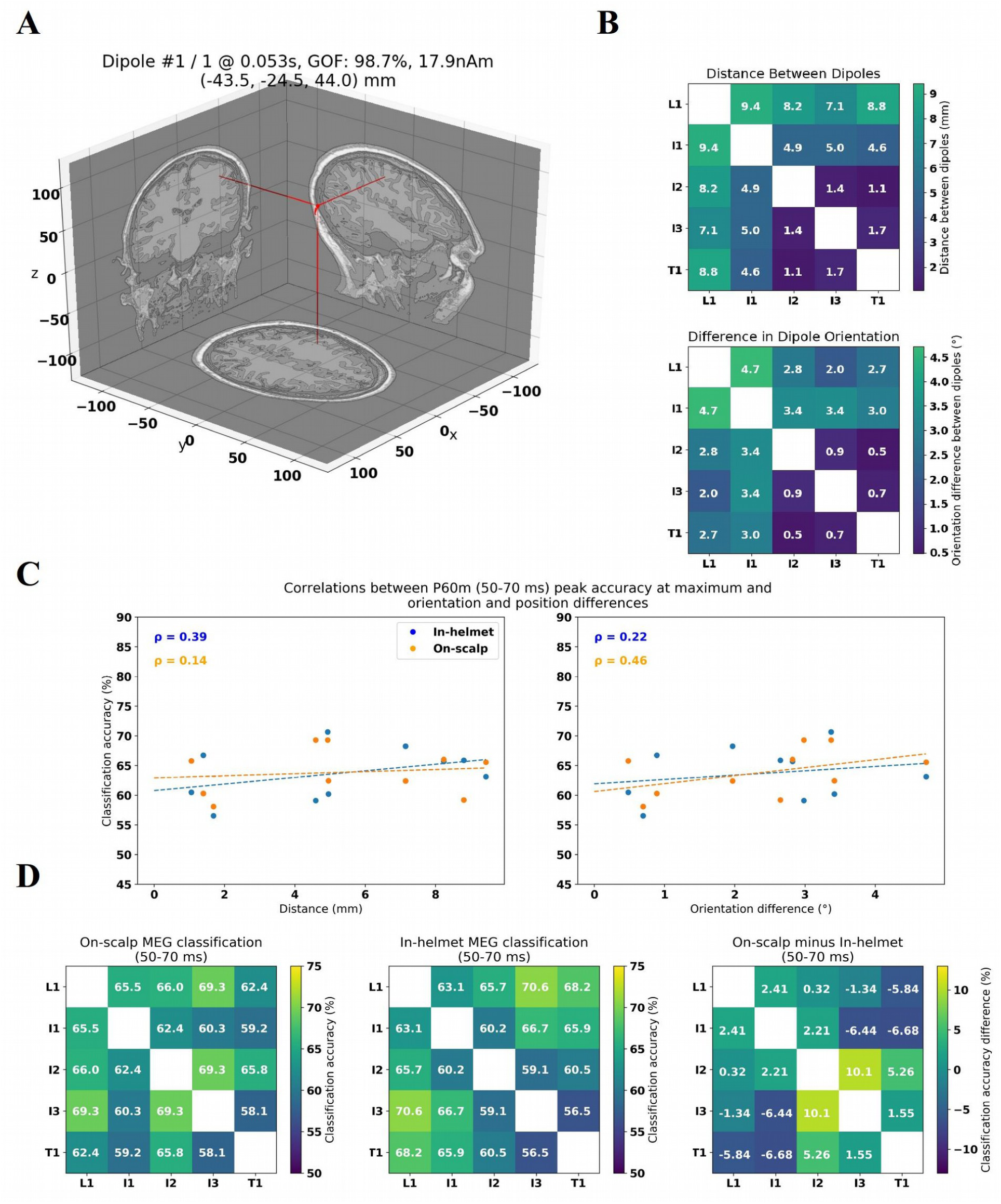
Dipole fit and peak classifications for Subject 4: **A:** Position and orientation of fitted dipole of I1. The dipole was fitted to contralateral SI. **B:** Distances and orientation differences between fitted dipoles. L1 is the one furthest away from all other dipoles and it also shows the greatest orientation difference compared to the others. **C:** Correlations between peak classification accuracy around P60m and difference in position and orientation for the maximum field position. **D:** Classification matrix of peak classification accuracy around P60m in the maximum recording for on-scalp and in-helmet magnetometers and the difference between them**-**See also Supplementary Figs. 2-6

### Classification – P60m (original hypothesis)

When plotting the peak classification accuracy in the window between 50 ms and 70 ms, the P60m, and plotting it against the differences in distance and orientations between the dipoles, positive correlations were found for both in-helmet and on-scalp data (Fig. 3D), thus supporting our hypothesis that phalanges with cortical representations further apart should be easier to classify than those with representations close to one another. However, we did not find evidence of on-scalp magnetometers classifying more accurately than the in-helmet magnetometers across the board (3D).

Unexpectedly, we found indications of classification accuracy rising more quickly for the on-scalp MEG as early as from 15 ms (Fig. 4-5). (See Supplementary Figs. 7-8., especially Subjects 2 and 4 for the early rise).

**Fig. 4:**
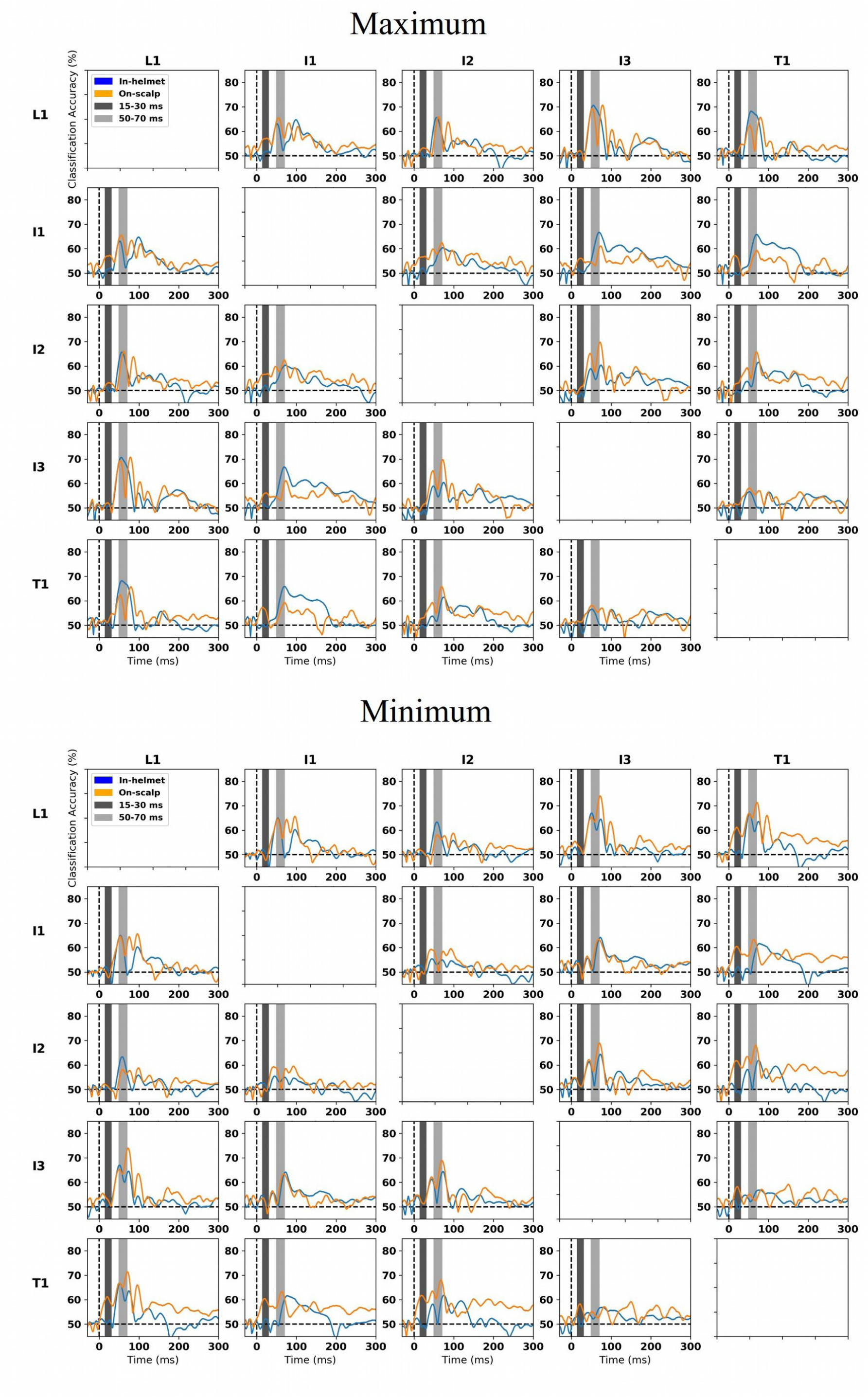
All-against-all classifications for the in-helmet and the on-scalp magnetometers above the maximum and minimum positions. The peak classification occurs around 50-70 ms (light grey) corresponding to the P60m. An interesting observation is that the on-scalp classification rises earlier than the in-helmet classification for the comparisons: L1-I1; L1-T1; I1-I2; I1-I3; I1-T1 on the maximum location (dark grey, 15-30 ms). For the minimum location, the same was the case for L1-T1, I1-T1, I2-T1, I3-T1. The 200 trials showing the highest variance were removed before classification. See also Supplementary Figs. 7-8

**Fig. 5:**
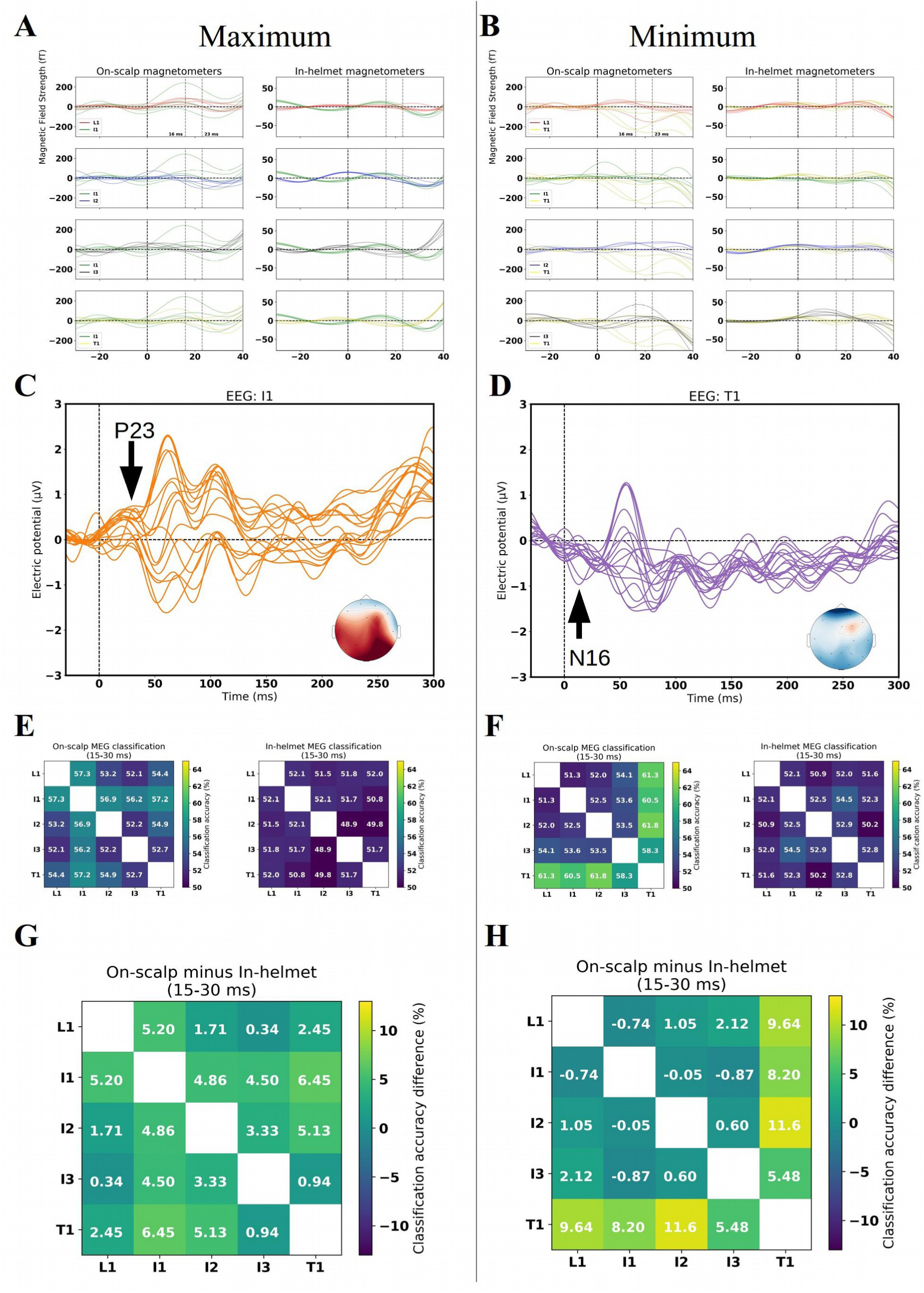
Early activity: **AB:** Zoomed-in view of the comparisons (Subject 4) where better classification accuracy was found for on-scalp than for in-helmet magnetometers. For the maximum position, comparing the two columns, it is seen visually that I1 differ more for on-scalp than for in-helmet magnetometers. For the minimum, the T1 is seen to differ more. **CD:** The EEG measurements for the I1 and T1 respectively showing early potentials, P23 and N16. The scalp topographies are for 23 ms and 16 ms. **E:** Classification for on-scalp and in-helmet classification for the maximum field for the early time period (15-30 ms). **F:** The same for the minimum field. **G:** The difference between on-scalp and in-helmet classification for the maximum field during the early time period. The comparisons including I1 show the biggest difference. **H:** The same for the minimum field – the comparison including T1 show the biggest difference. See also Supplementary Figs. 9-12.

### Classification – early activity (unexpected results)

To better understand the differences in early classification accuracy, we zoomed in on the early part of the ERFs and the classification accuracy (Fig. 5 & Supplementary Figs. 9-12). In the maximum recording, we found, visually, that the ERF picked up by one of the on-scalp magnetometers for I1 differed clearly from the others in the time range of 15 ms to 30 ms. In the minimum recording, the ERFs for the T1 picked up by several on-scalp magnetometers differed clearly in the same time range (Fig. 5AB). In the EEG, we found ERPs in this very time range (Fig. 5 C-F), specifically the N16 and P23 (Tsuji and Murai, 1986; Yamada et al., 1984).

### Grand averages

To investigate where the classifications might differ, we calculated the *t*-values (*df*=3) (Student, 1908) for the contrast between in-helmet and on-scalp classification for each time sample. The results are not corrected for multiple comparisons. We found that the P60m gave rise to significantly better classification for the in-helmet magnetometers compared to the on-scalp magnetometers for many of the comparisons (Fig. 6A). The grand average however showed the same early (significant) effect (Fig. 6AB) found in Subject 4 (Fig. 5). See Supplementary Fig. 13 for the corresponding figure for the minimum position.

**Fig. 6:**
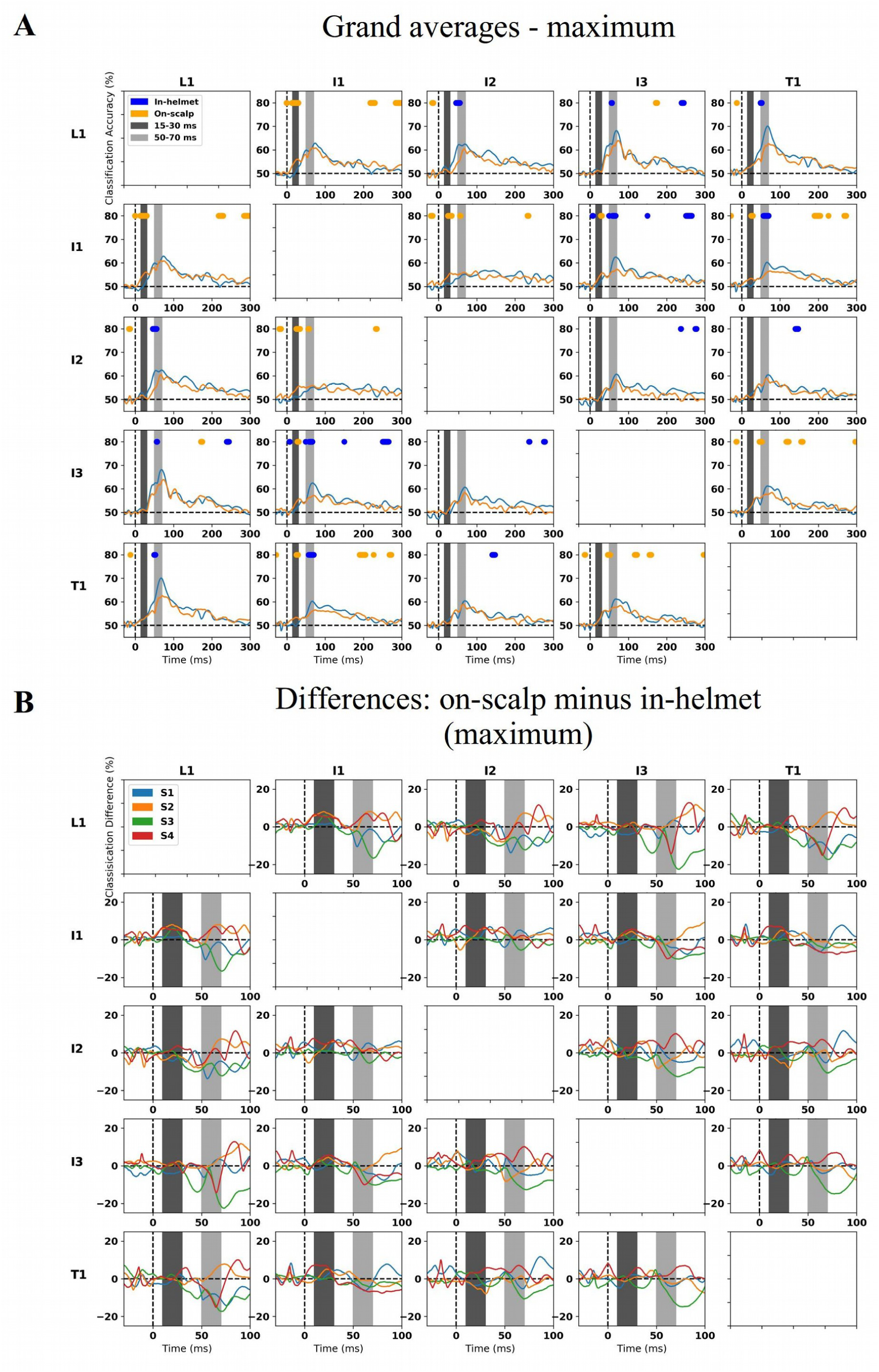
All subjects for the maximum position. **A:** Grand averages of the decoding performance across all four subjects with dots indicating where classifications differed significantly (t-distribution with df=3) between on-scalp and in-helmet magnetometers. Orange dots indicate that on-scalp was better and blue dots that in-helmet was better. Looking at the early time interval (15-30 ms), it is seen that L1-I1; I1-I2; I1-I3 and I1-T1 all have better classification for the on-scalp measurements. For the P60m component, it is seen that in-helmet measurements in general (significantly) outperforms on-scalp magnetometers. **B:** Differences between on-scalp and in-helmet classification. Especially, in the I1 row it is clear that the subject focused on (Subject 4) is not the only subject showing early effects for on-scalp magnetometers. See Supplementary Fig. 13 for the minimum position.

## Discussion

In this study, we set out at comparing spatio-temporal resolution between on-scalp and in-helmet MEG by assessing the discrimination accuracy for activity patterns resulting from stimulating five different phalanges of the right hand. Because of proximity and spatial resolution differences between on-scalp and in-helmet MEG, we hypothesized that on-scalp MEG would allow for better classification performance as compared to state-of-the-art MEG.

Our results show that the P60m response component in both on-scalp MEG and in-helmet MEG could be used to successfully discriminate between all the activations patterns of the somatosensory cortex, both between fingers (I1, L1 and T1), and within fingers (I1-I3) (Figs. 3-4, Supplementary Figs. 5-6). As initially hypothesized, these results also confirmed that classification accuracy correlates positively with greater distances between estimated sources and also with greater differences in orientations between contrasted sources (Fig. 3B, Supplementary Figs. 3-4).

However, our initial hypothesis that on-scalp MEG would allow for better classification performance as compared to state-of-the-art MEG around the P60m was not confirmed. If anything, it was the other way around with in-helmet magnetometers outperforming on-scalp magnetometers. We discuss the possible reasons to and implications of this in a separate section below.

### Early activity (15-30 ms)

The comparison of on-scalp and in-helmet discrimination accuracy however also generated an incidental finding, where on-scalp MEG recordings resulted in a superior classification performance as compared to in-helmet MEG. These findings stem from the time period *before* the rise of the P60m, in the time range of 15-30 ms (Figs. 4&6) and were linked to the N16 and P23 components found in EEG recordings. Furthermore, these responses were seen only in the on-scalp MEG and not in the in-helmet MEG data (Fig. 5). This was particularly clear in two out of the four subjects (Supplementary Figs. 9-10) while the advantage in classification performance for on-scalp over in-helmet MEG was also shown to be overall statistically significant (Fig. 6).

### Thalamo-cortical radiation

In all our subjects, we observed a peak around 16 ms (Figs. 5AB & Supplementary Figs. 11-12). Several studies have provided evidence that the thalamo-cortical radiation between thalamus and S1 can be picked up by EEG (Buchner et al., 1995; Gobbelé et al., 2004; Götz et al., 2014). It might therefore also be theoretically possible with MEG even though it is complicated by these sources being deep and extending towards the centre of the brain. Some evidence has been found of the thalamo-cortical radiation being possible to reconstruct with dipole fitting using conventional MEG (Kimura et al., 2008; Papadelis et al., 2012). These studies however did not find clear evidence of a component in the sensor-level data, as we did, but based their source reconstruction on *a priori* information from functional Magnetic Resonance Imaging (fMRI) and EEG studies.

### The quasi-radial nature of P23

The literature shows that the source underlying the P23 is radially oriented (Deiber et al., 1986). Radially oriented sources are theoretically invisible to MEG, but visible to EEG. The theoretical invisibility of radially oriented sources only holds though, in the ideal case where the head is assumed to be perfectly spherical. De Munck and Daffertshofer (2012) discuss that radial sources are in practice at most quasi-radial, in that they are not completely invisible, but that they are 5 to 50 times weaker than fully tangential sources (Fuchs et al., 1998). A straightforward explanation for why on-scalp MEG may pick up the P23m while in-helmet MEG does not, is thus that the signal is so weak that it deteriorates beyond detectability before reaching the in-helmet magnetometers (i.e. it is a question of SNR due to differences in proximity). From the present results alone, we cannot conclude that the difference in proximity is the sole cause of the observed difference. An important difference between the on-scalp and in-helmet measurements performed here is also that the pickup coils differ in size, 9.2 mm × 8.7 mm for on-scalp MEG versus 21 mm × 21 mm for in-helmet. Interestingly, Haueisen et al. (2012) were able to detect the P23m using low-*T*_*c*_ SQUIDs with a pickup coil size of 8 mm × 8 mm. They found that the P23m was more spatially focal and sharper than the N20m. The P23 has been localized to Brodmann Area 1 (Buchner et al., 1994), which is very close to Brodmann Area 3 ∼1-2 cm, where the dipoles were localized to for the P60m in the present study Fig. 3).

If the magnetic field related to the quasi-radial source is sharp and focal as reported by Haueisen et al. (2012), and not wide and distributed, a large coil may average away the signal due to the field lines closing on themselves within a large coil. The advantage for on-scalp magnetometers during the early period may thus be due to the finer spatial sampling with on-scalp magnetometers picking up the focal weak quasi-radial signals that would be averaged away in coils that sample a larger area.

It is thus not unlikely that some of the sensors of the present array, which are spaced ∼11 mm apart (Fig. 1) happened to be placed at a position where the P23m can be picked up. Based on the present recording, however, we cannot ascertain whether the increase in classification accuracy for on-scalp relative to in-helmet magnetometers is due to differences in coil size or differences in SNR (or a combination of the two). It should also be noted that there is a trade-off between the size of a pickup coil and its sensitivity/specificity. That is, larger coils are more sensitive, but less specific, and vice versa for smaller coils. The bigger the pickup coil, the more sensitive it is to a more uniform field, but the less specific it is, i.e. it picks up more activity, but its spatial resolution decreases. Roughly speaking, when sensors are far away, you need to weigh sensitivity over specificity, and when sensors are close by, you can relax the sensitivity in favour of better specificity, i.e. spatial resolution. Thus, in terms of making a comparison where it can be estimated how much each of the two factors, coil size and SNR, contribute to the increased classification accuracy for the on-scalp magnetometers, it is not meaningful to build on-scalp magnetometers with an area the size of the in-helmet magnetometers used here (21 mm × 21 mm), because the decrease in specificity would counteract the idea of moving closer to the scalp (i.e. achieving increased spatial resolution). Nonetheless, disentangling the effects of coil size and SNR is important, but remains to be a job guided primarily by simulation studies (Iivanainen et al., 2017).

### Further discussion of P60m classification

The motivation for the hypothesis that on-scalp classification would result in better classification than in-helmet classification was the improved sensor-to-brain proximity, and the associated gain in SNR. A feature of the data that we could not fully evaluate beforehand here is whether the sensor array would be large enough to pick up both the common activations between phalanges *and* the unique activations for each phalange. To investigate whether this was the case, we evaluated the overlap of activations for each phalange at 60 ms (Fig. 7). See Supplementary Fig. 14 for all subjects.

**Fig. 7:**
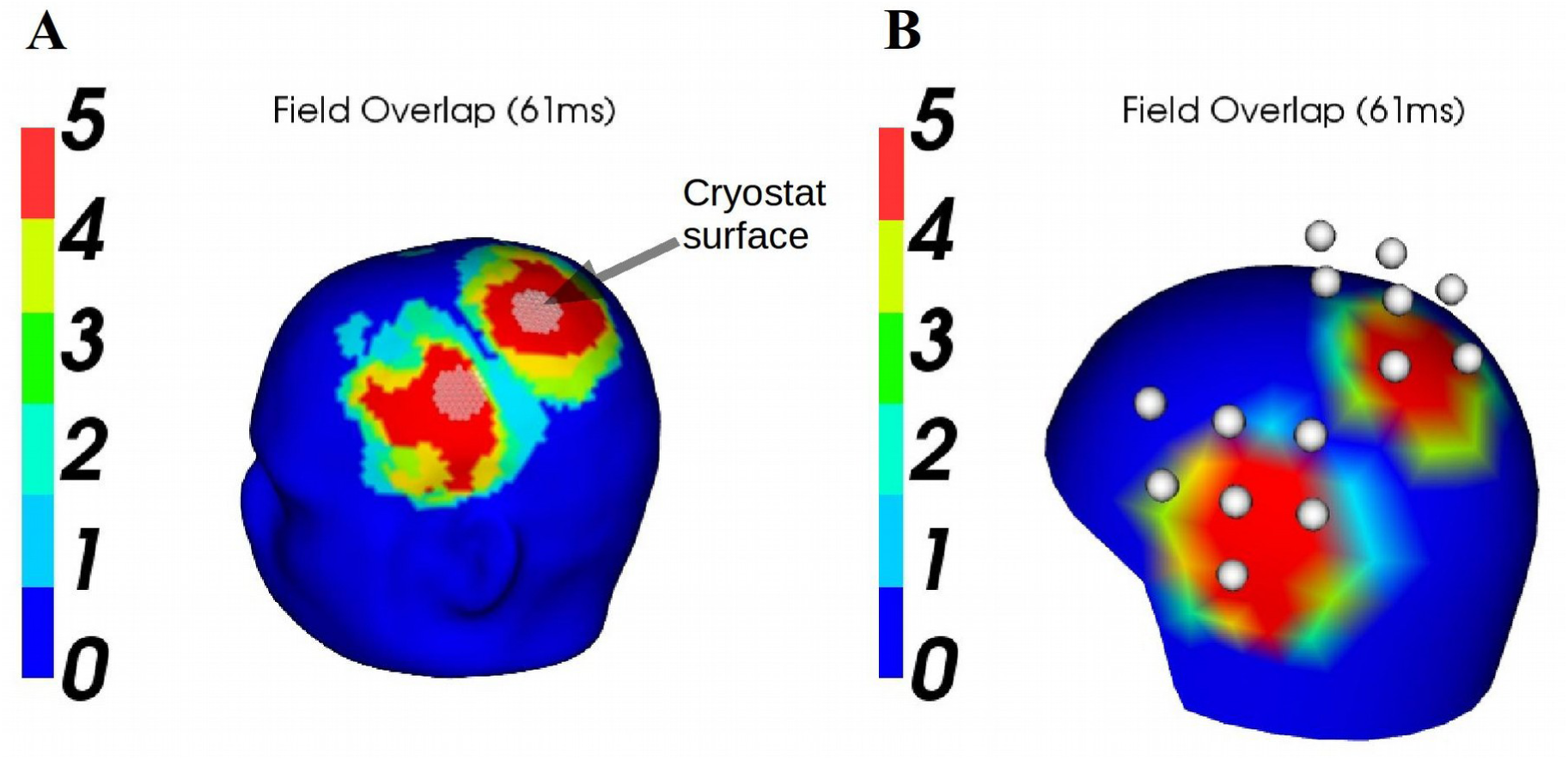
Relation between spread of field and sensor positions for Subject 4. **A:** The overlap of fields generated by the dipole models for each phalange at 61 ms is illustrated. The lower and upper fifth percentiles of magnetic field strength on the head were calculated for each of these dipole models. The colour map indicates where these percentiles overlap. The white dots indicate the surface of the cryostat. It is seen that the cryostat was placed where the fields overlap either completely or at least four out of five do. **B:** The same plot for the in-helmet recording, but the white dots indicate the actual positions of the seven sensors used for classification. Note that the overlap between the fields here is visibly different from the **A** panel. See also Supplementary Fig. 14.

These plots of overlapping fields indicated that the on-scalp MEG sensor array mostly picked up signal from areas that overlap either completely or greatly in terms of signal strength (Fig. 7A). For the in-helmet MEG sensors used, results show less overlap between sensors (Fig. 7B).

Consequently, it was the case that the signal at each measurement point was more homogeneous within the area covered by the on-scalp MEG than by the in-helmet MEG sensors indicating that we had placed the on-scalp MEG at sub-optimal positions.

### Future directions

The findings above indicate that to fully evaluate our initial hypothesis regarding spatio-temporal resolution for the P60m, it is necessary to repeat this study with an on-scalp sensor array that covers both field commonalities and field differences for the involved between-phalange comparisons; or in the absence of a larger sensor array at least record positions around the extrema to provide full coverage of the fields.

In a similar way, it would be of great interest to evaluate the early activity measured here, N16m and P23m, more thoroughly with more dedicated on-scalp experiments. First, the component could be localized, preferably using high-density EEG. Then, the corresponding MEG field could be projected using anatomically accurate volume conductors, and then place on-scalp sensors accordingly. Another option, which is possible with extensive sensor on-scalp MEG sensor coverage such as already possible with OPMs, the whole somatosensory cortex may be covered at one go. One could of course also use a small sensor array as here but perform recordings at several places above the somatosensory cortex so to accumulate coverage of a larger area.

## Conclusions

In our comparison of spatio-temporal resolution between on-scalp and in-helmet MEG, we did not find the hypothesized improved classification for on-scalp MEG compared to in-helmet MEG around the P60m. We suggest that an important reason for this unexpected finding is that the fields associated with the five phalanges overlap to a high degree, and that the on-scalp sensors cover too small an area to discern the differences between these fields (Fig. 7). In an unexpected finding, we however show that the on-scalp MEG classified the phalanges better than the in-helmet MEG during an early time period (15-30 ms), which may reflect the thalamo-cortical radiation from thalamus to S1 and/or the quasi-radial P23m. We propose that these findings stem from the improved signal gain and/or the increase in spatial sensor resolution using on-scalp MEG.

## Acknowledgements

This work was funded by Knut och Alice Wallenbergs Stiftelse (KAW2014.0102), the Swedish Research Council (2017-00680) and the Swedish Childhood Cancer Foundation (MT2018-0020).

Data for this study was collected at NatMEG, the National infrastructure for Magnetoencephalography, Karolinska Institutet, Sweden. The NatMEG facility is supported by Knut & Alice Wallenberg (KAW2011.0207)

We also acknowledge the contribution of Veikko Jousmäki, Aalto University, Finland for building the custom stimulation system.

