## Supplementary Figures 1-14 for "On-scalp MEG SQUIDs are sensitive to early somatosensory activity unseen by conventional MEG"

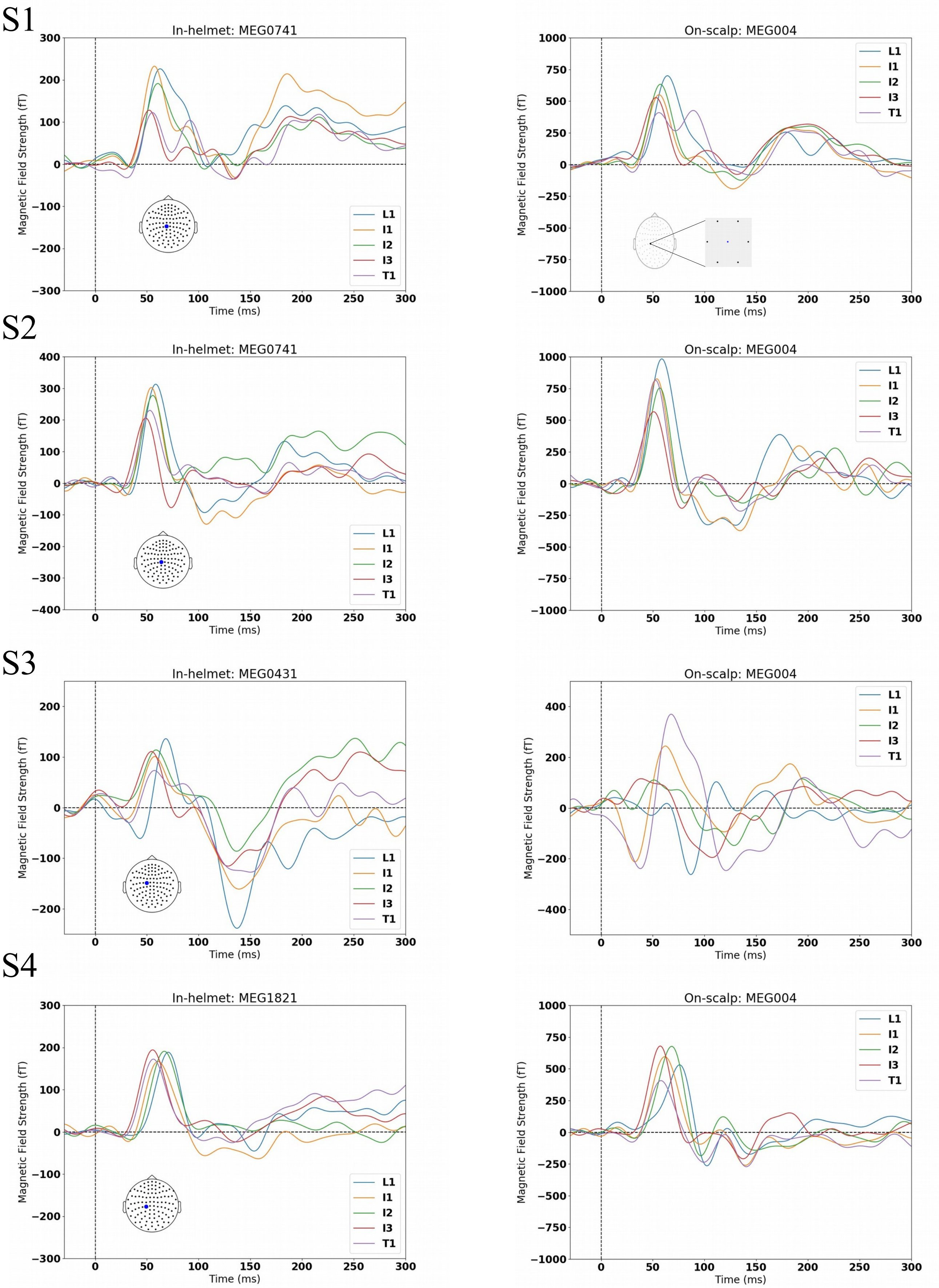


*Supplementary Fig. 1: The maximally responding in-helmet magnetometer alongside the centre on-scalp magnetometer are shown for each subject. Note that the P60m of Subject 3 is not clearly defined for all phalanges for the on-scalp measurement. Compare with Fig. 2. in the manuscript.*

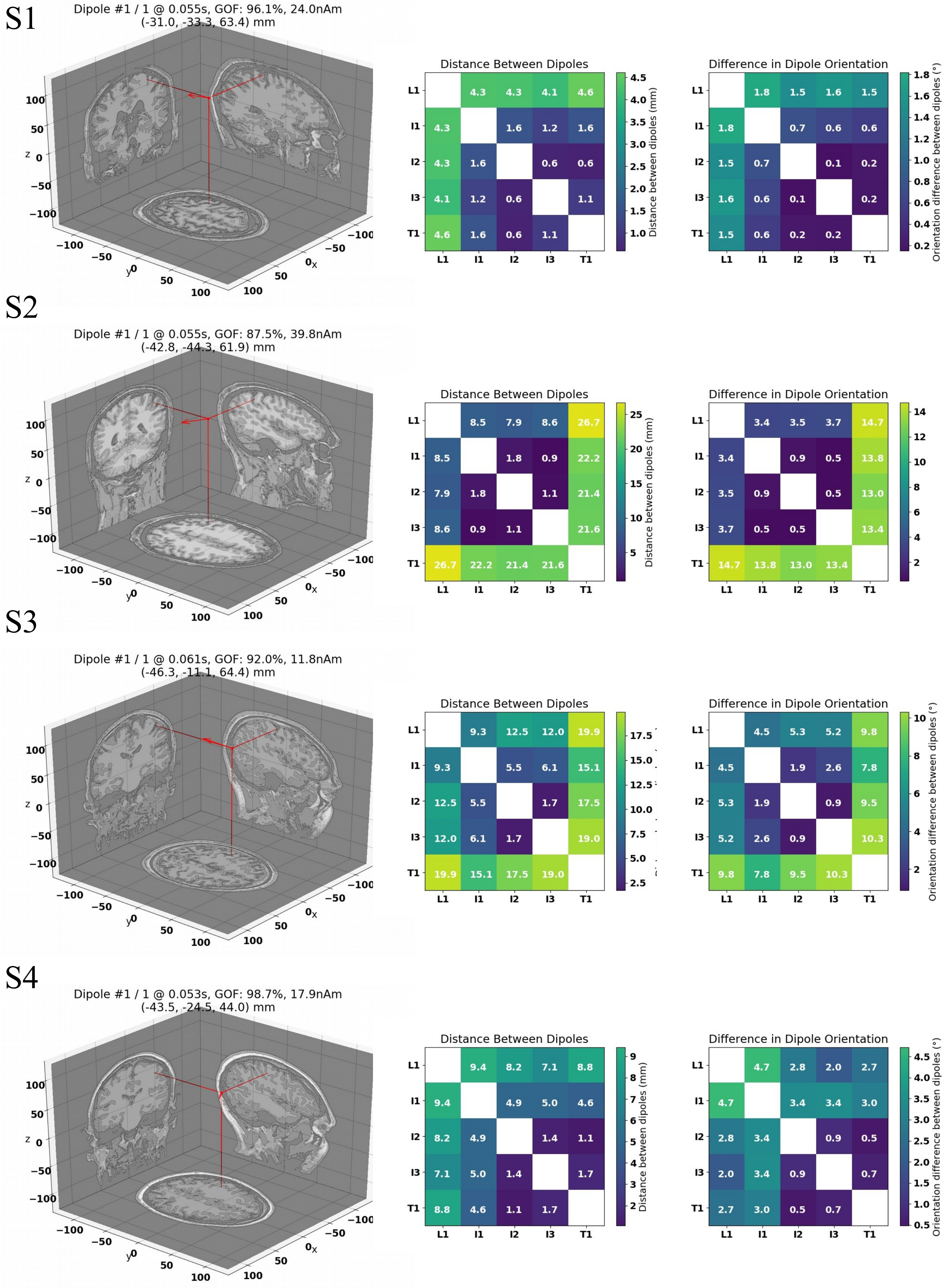


*Supplementary Fig. 2: Dipole fits for position I1 for each subject alongside matrices showing the differences in distance and orientation. Compare with Fig. 3. in the manuscript.*

S1

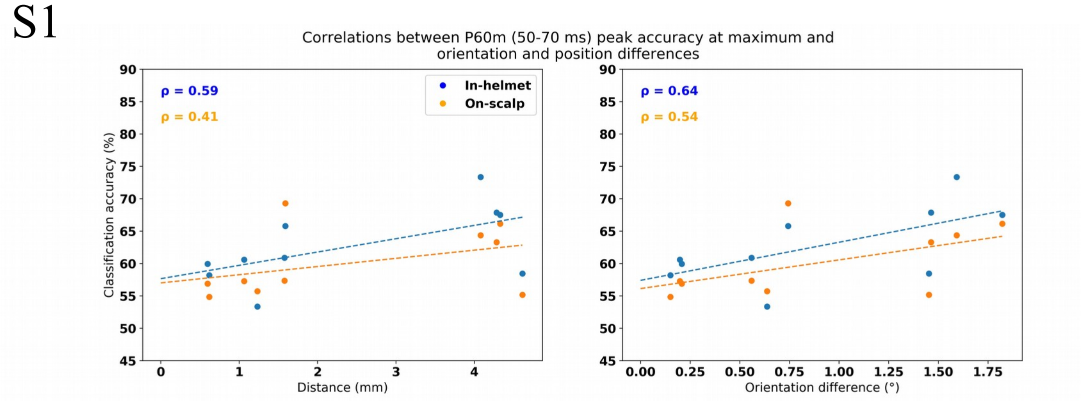


S2

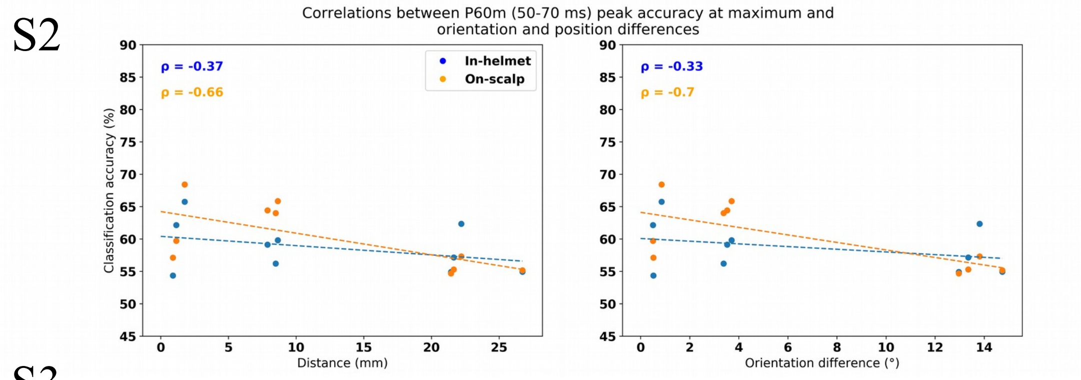


S3

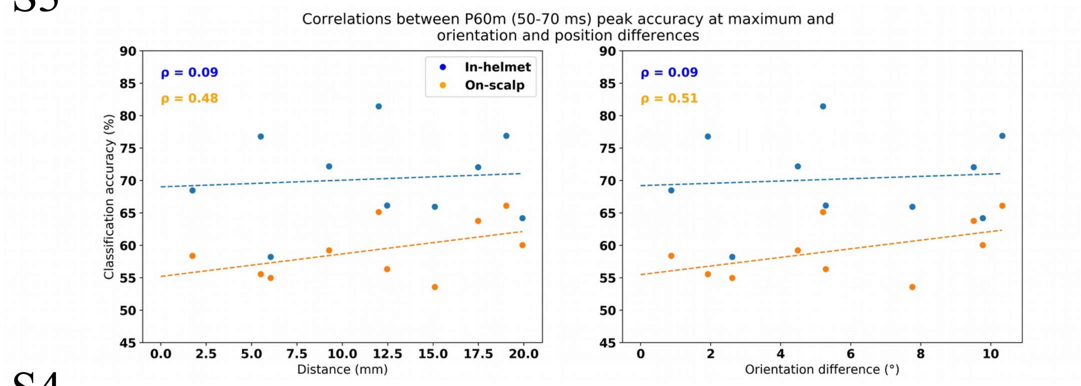


S4

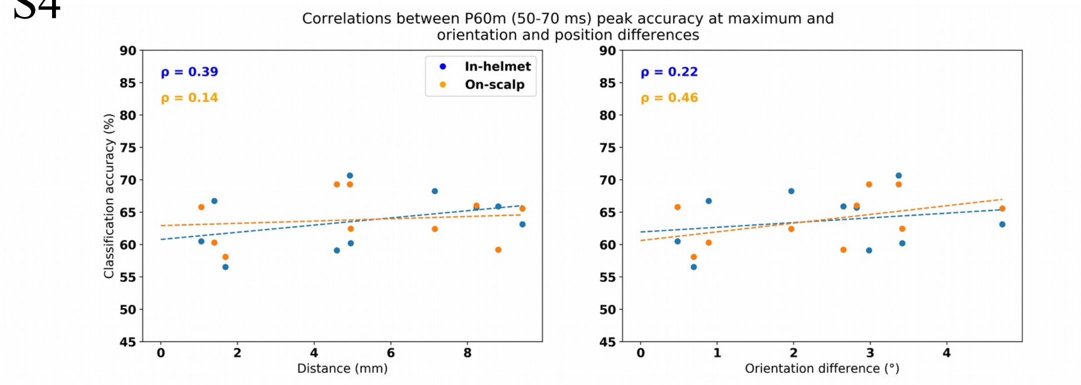


*Supplementary Fig. 3: Correlations between P60m peak accuracy and orientation and distance at the maximum position for each subject. Subject 2 unexpectedly showed negative relationships. Compare with Fig. 3. in the manuscript.*

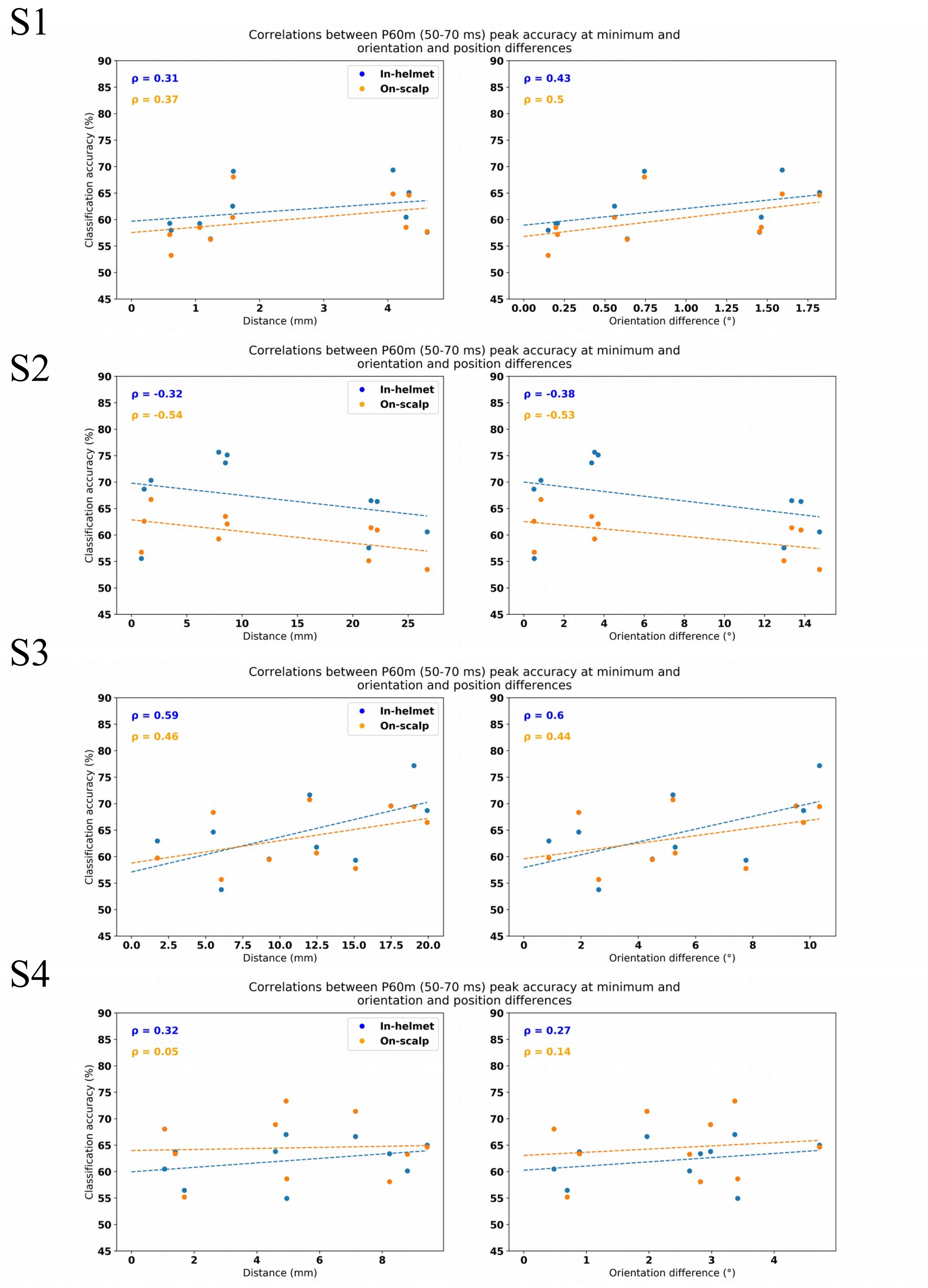


*Supplementary Fig. 4: Correlations between P60m peak accuracy and orientation and distance at the minimum position for each subject. Subject 2 unexpectedly showed negative relationships. Compare with Fig. 3. in the manuscript.*

### Maximum

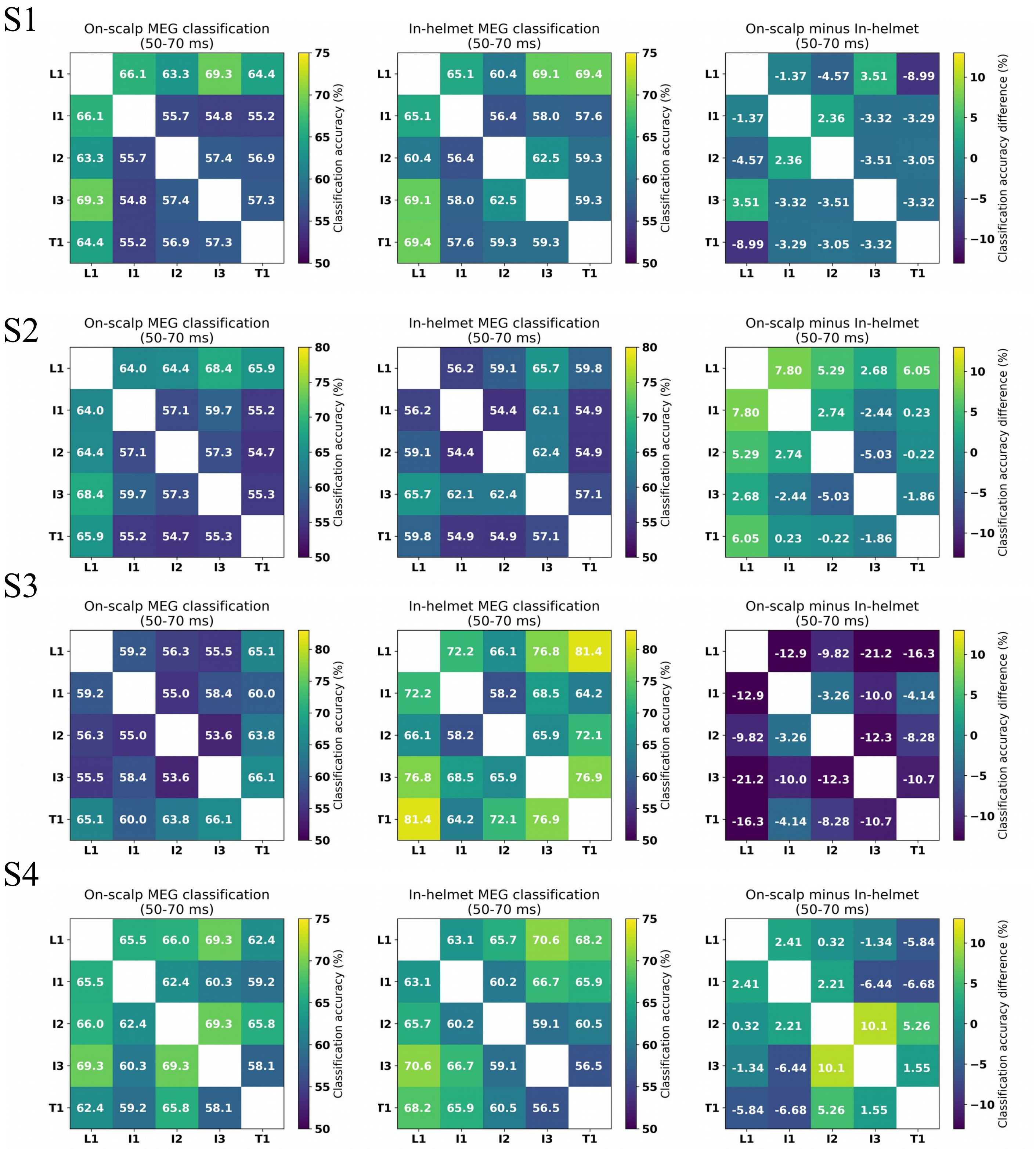


*Supplementary Fig. 5: Peak classification during the P60m period (50-70 ms) for the maximum position. Note that Subject 3 stands out showing better classification across the board for in-helmet classification, whereas it is more variable for the remaining subjects. Subject 2, for example, in general shows an advantage for on-scalp classification. Compare with Fig. 3. in the manuscript.*

### Minimum

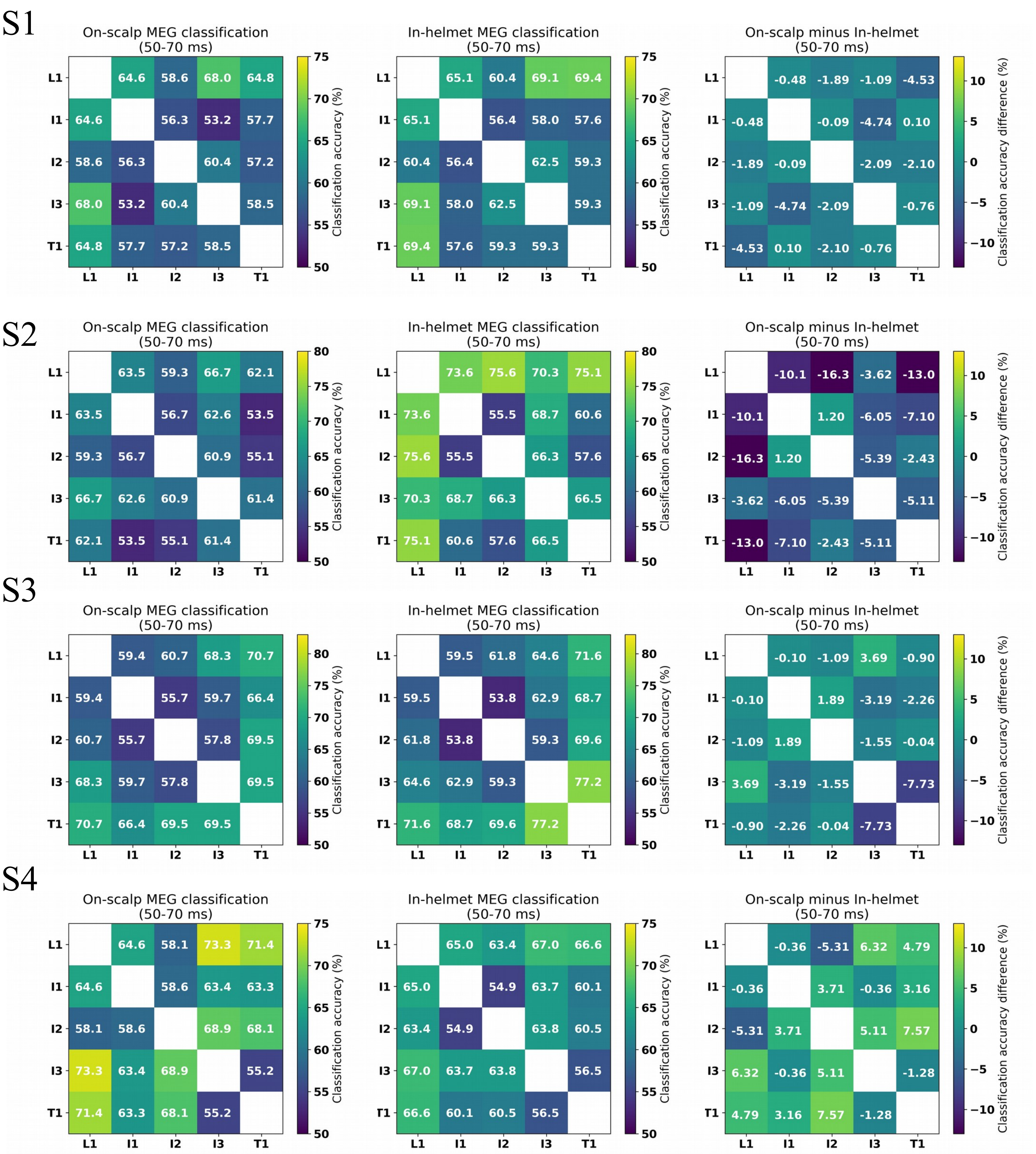


*Supplementary Fig. 6: Peak classification during the P60m period (50-70 ms) for the minimum position. Compare with Fig. 3. in the manuscript.*

S1

### Maximum

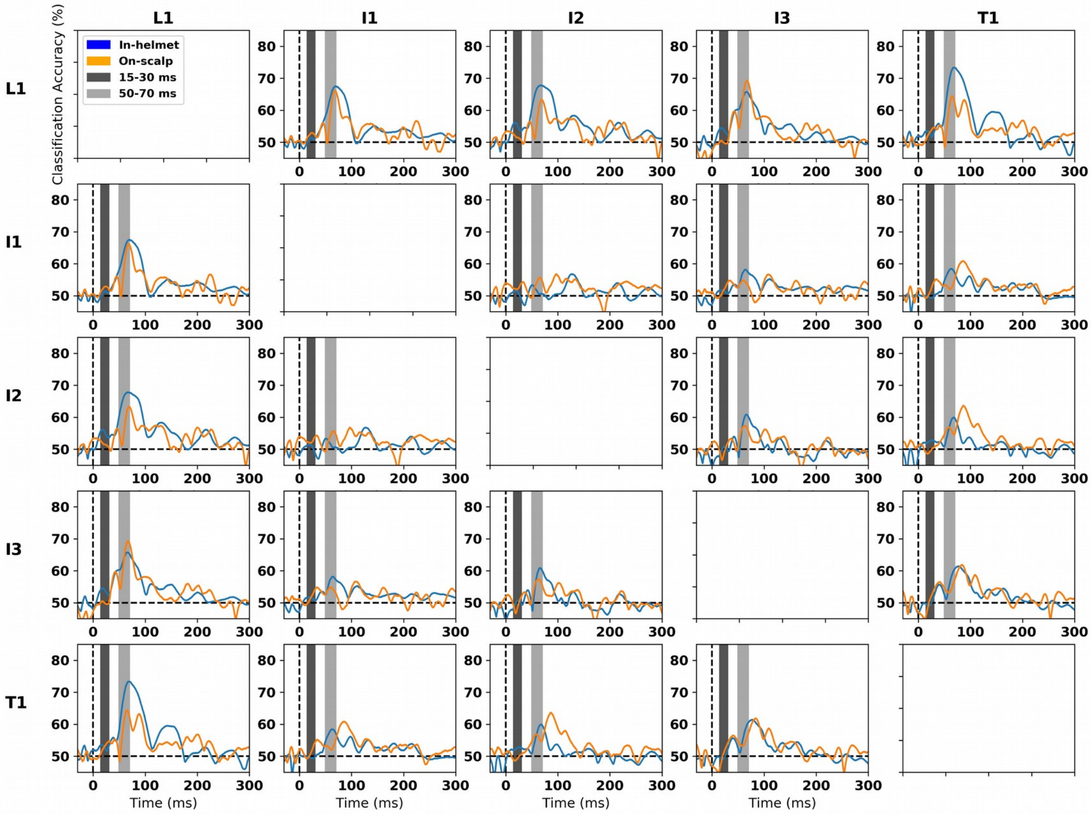


S2

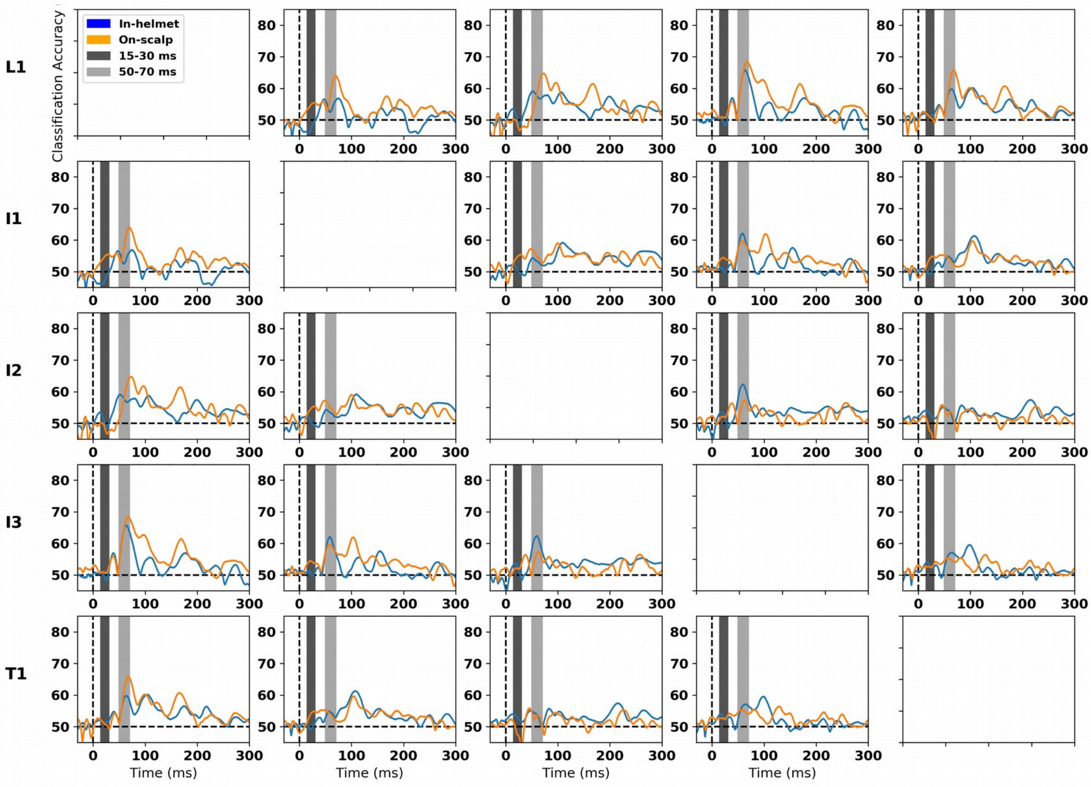


Continued on next page

S3 Maximum Continued from above

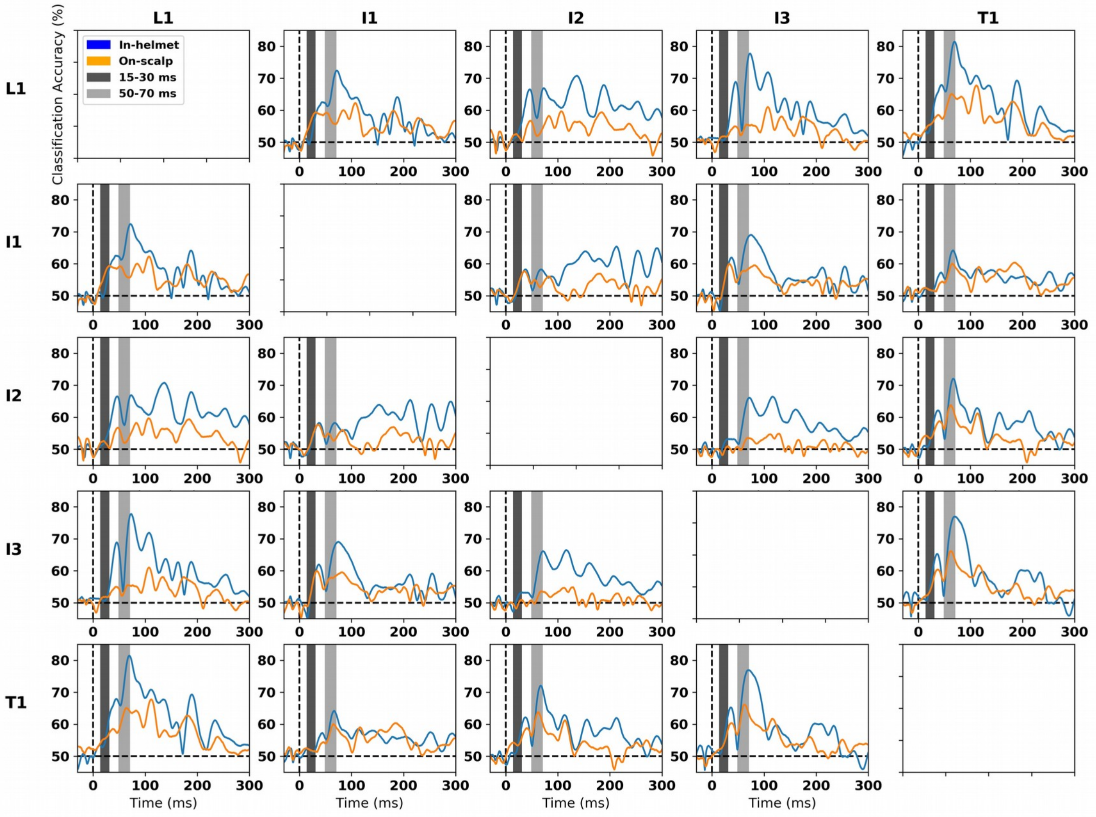


S4

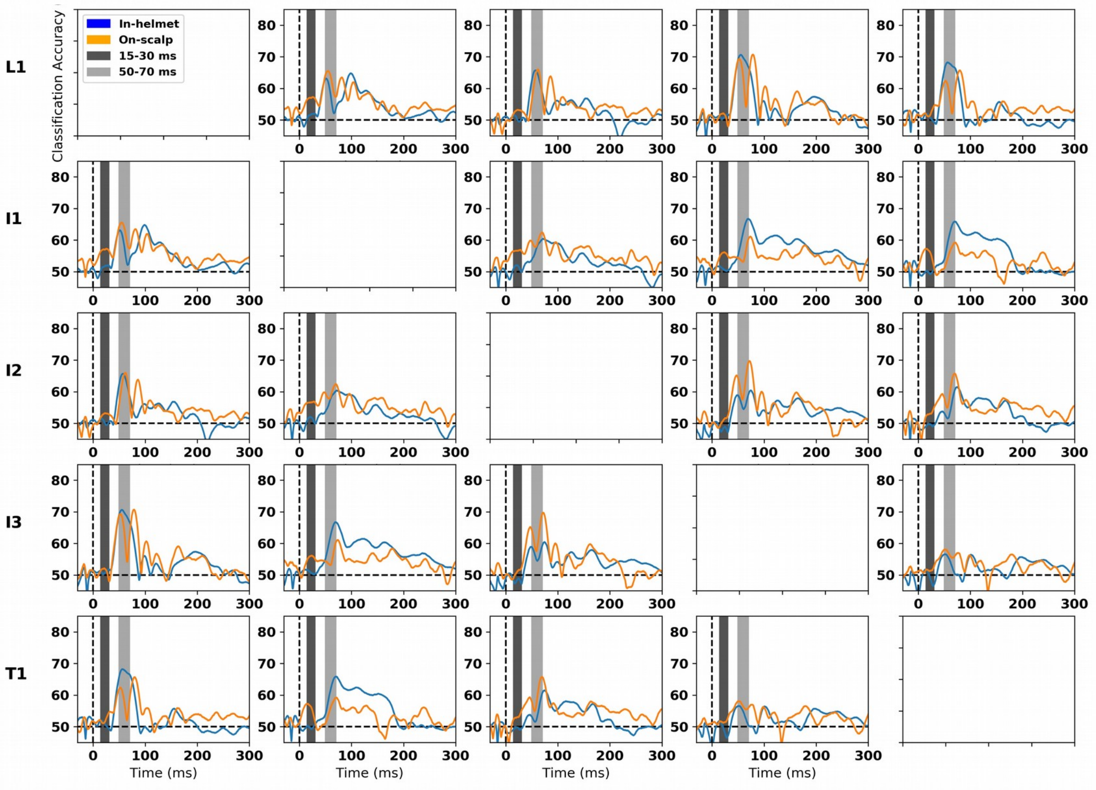


*Supplementary Fig. 7: Classification time courses for the maximum position for all subjects. Note again the individual differences also observed in Supplementary Fig. 5. Subject 3 stands out showing better classification across the board for in-helmet classification, whereas it is more variable for the remaining subjects. Subject 2, for example, in general shows an advantage for on-scalp classification even during the P60m. Subjects 2 and 4 show the strongest early (15-30 ms) differences. Compare with Fig. 4. in the manuscript.*

S1

### Minimum

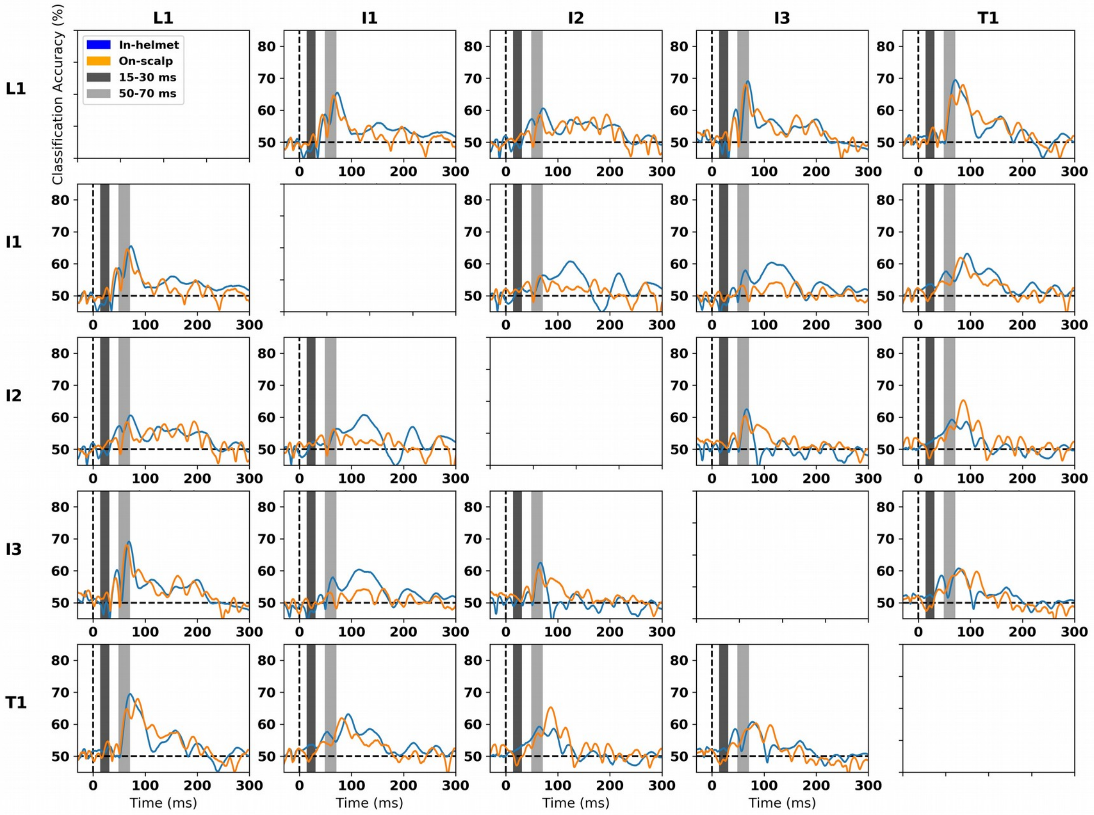


S2

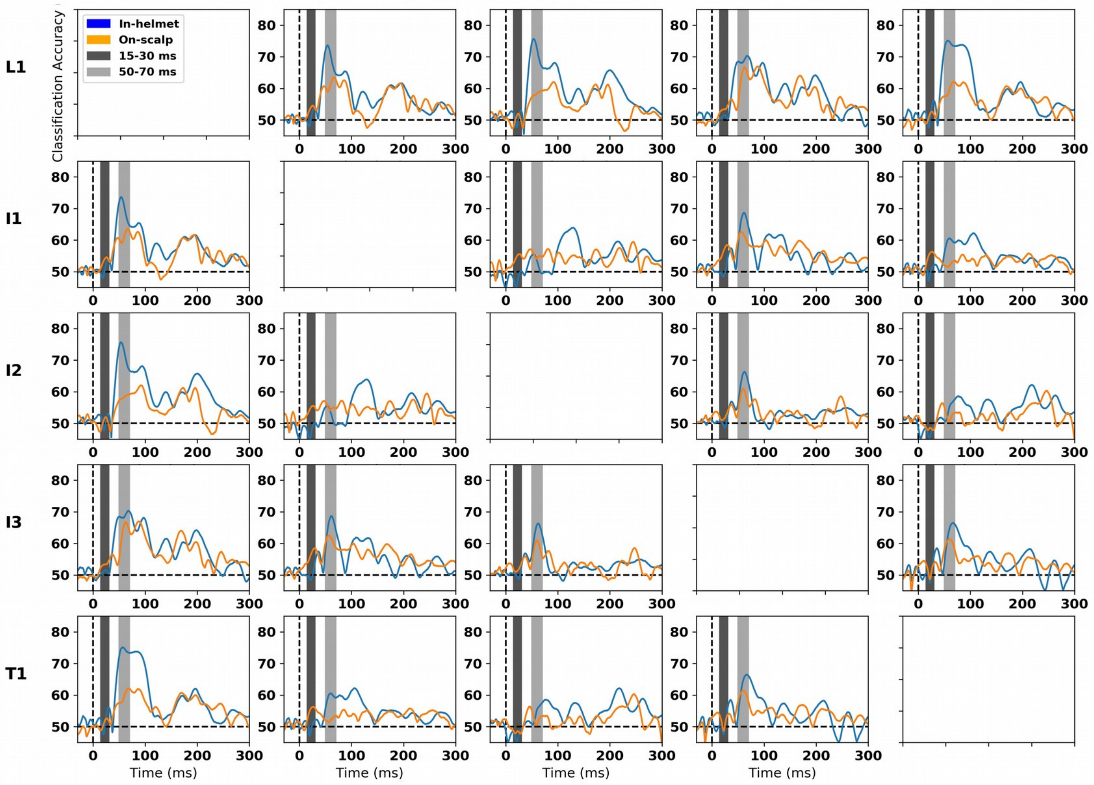


Continued on next page

S3

Minimum

Continued from above

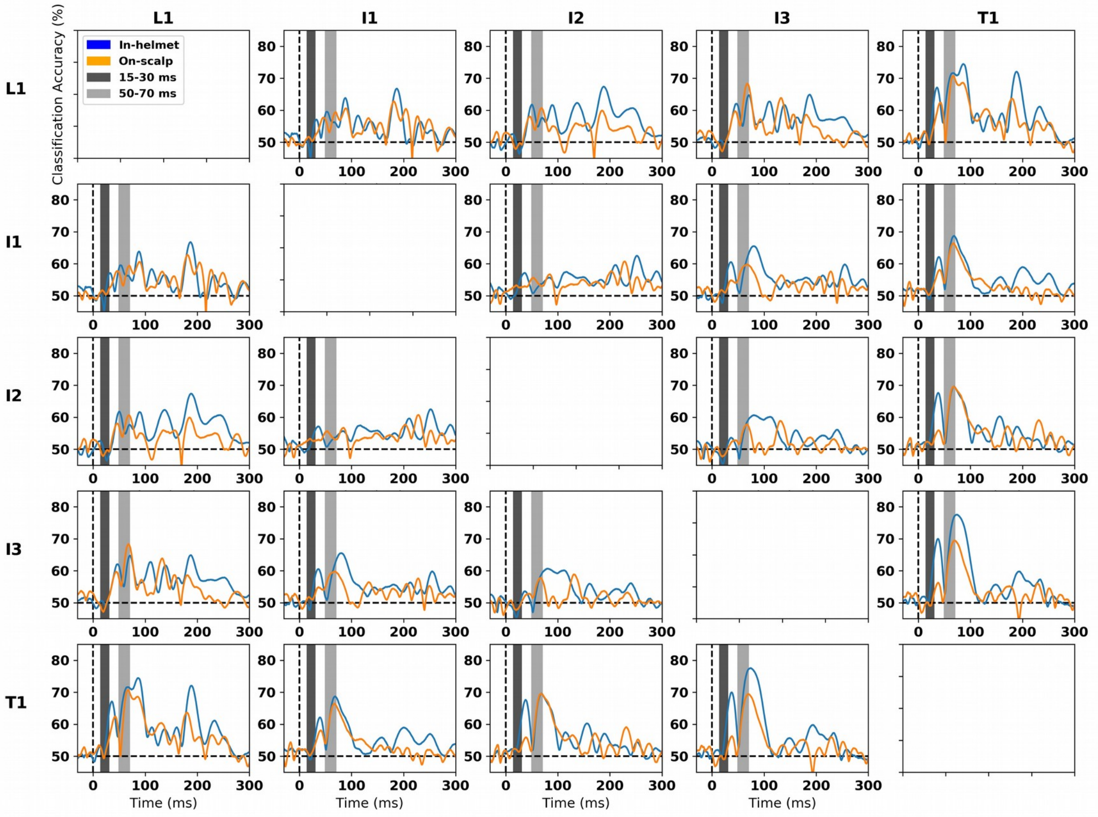


S4

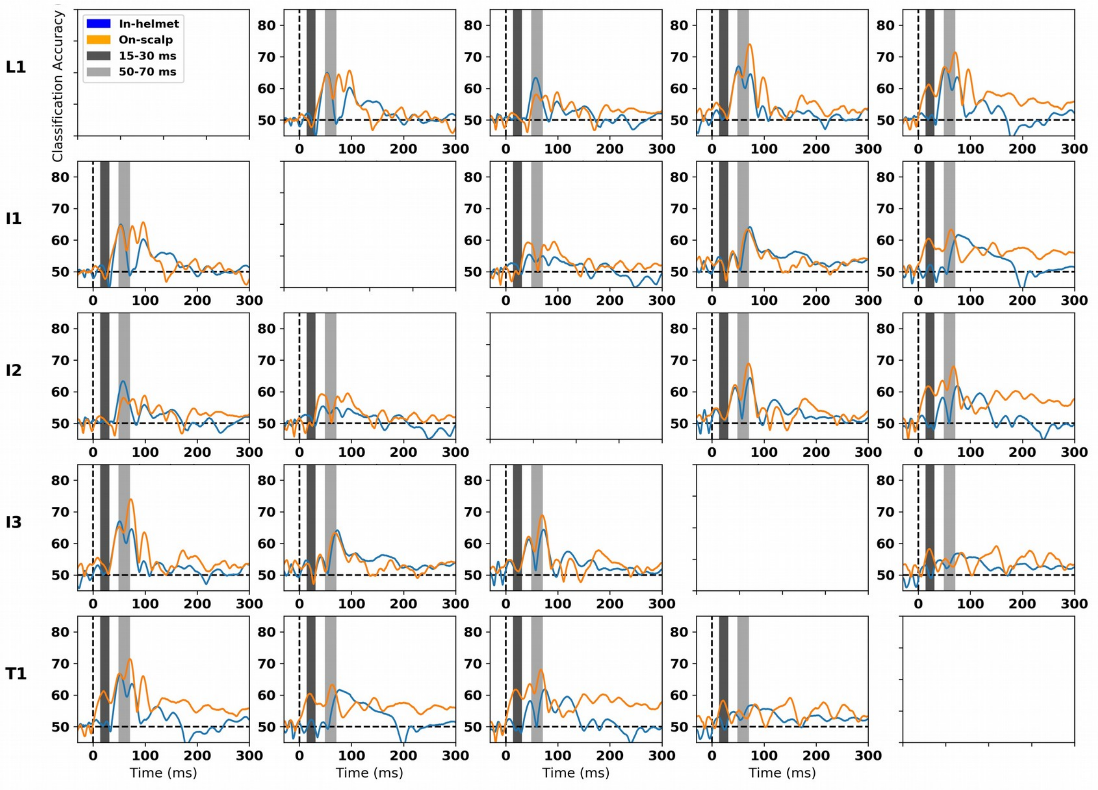


*Supplementary Fig. 8: Classification time courses for the minimum position for all subjects. Note again the individual differences also observed in Supplementary Fig. 4. Subject 2 stands out showing better classification across the board for in-helmet classification, whereas it is more variable for the remaining subjects. Subject 4, for example, in general shows an advantage for on-scalp classification even during the P60m. Subjects 2 and 4 show the strongest early differences (15-30 ms) for respectively I1 and T1. Compare with Fig. 4. in the manuscript.*

### Maximum

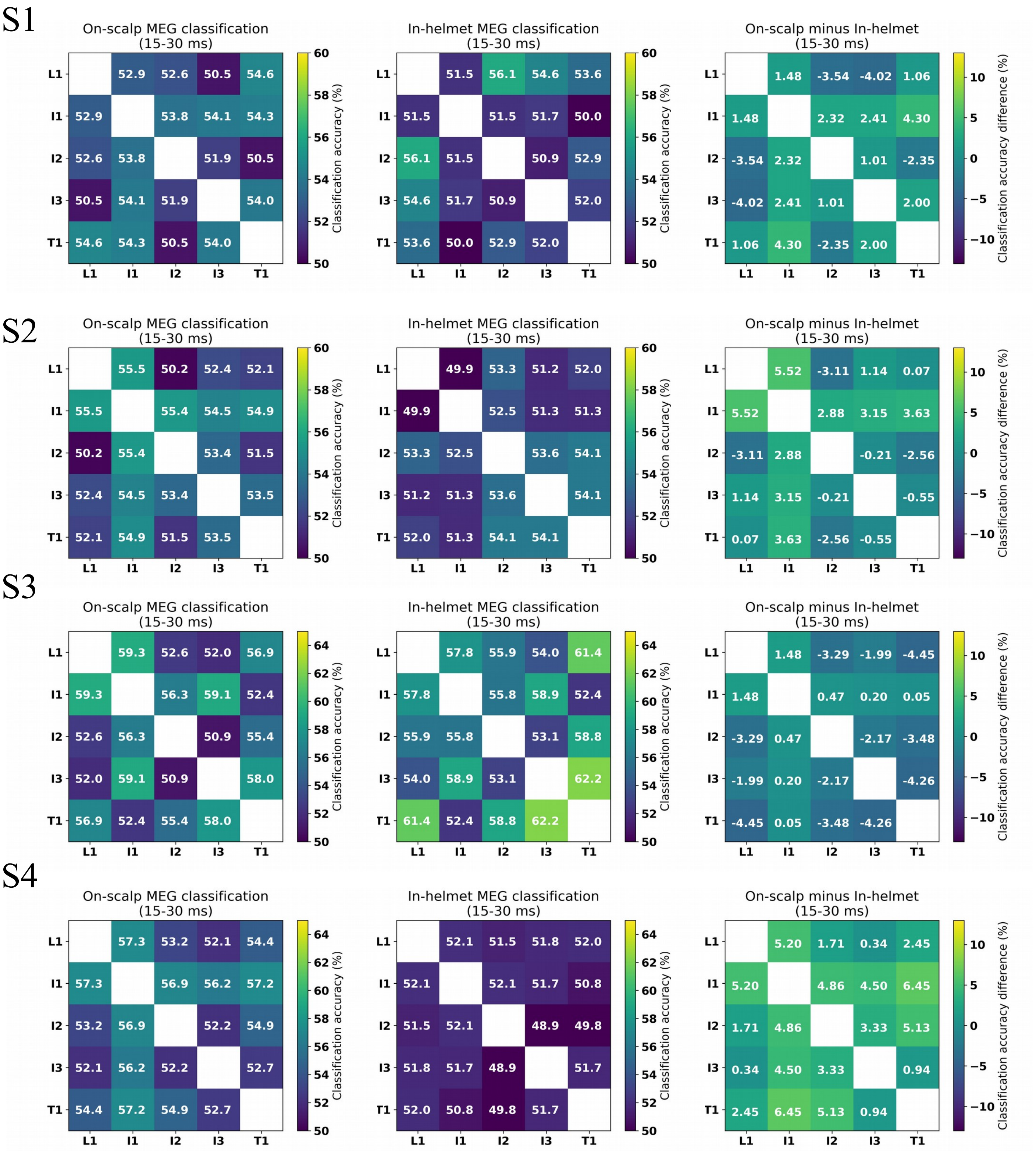


*Supplementary Fig. 9: Peak classification during the early time period (15-30 ms) for the maximum position. Note that Subjects 2 and 4 stand out showing better classification for I1 for on-scalp classification. Compare with Fig. 5 in the manuscript.*

### Minimum

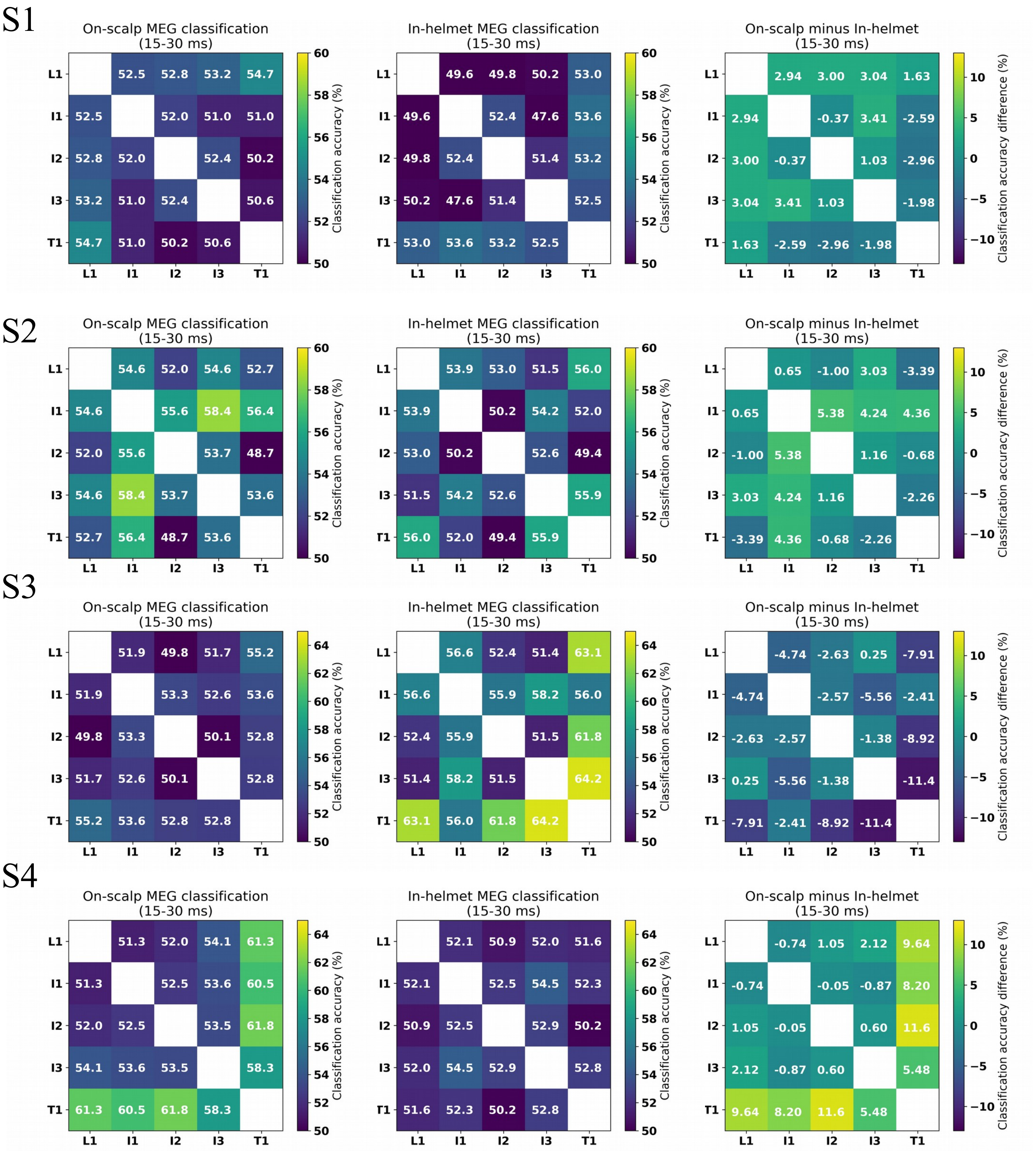


*Supplementary Fig. 10: Peak classification during the early time period (15-30 ms) for the minimum position. Note that Subjects 2 and 4 stand out showing better classification for I1 and T1 respectively for on-scalp classification. Compare with Fig. 5 in the manuscript.*

### Maximum

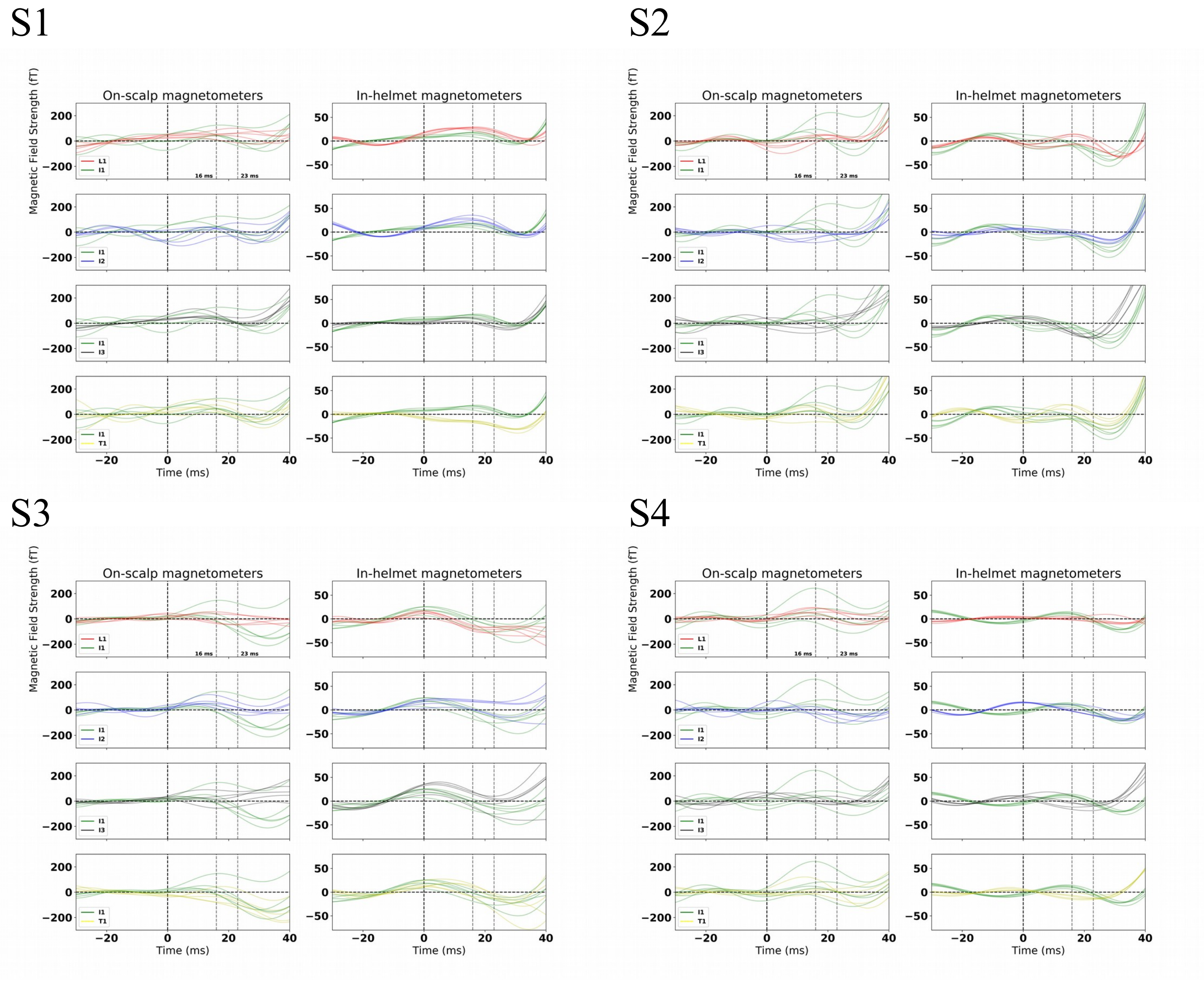


*Supplementary Fig. 11: Zoomed in view of the evoked responses (-30 to 40 ms) for the maximum position for the comparison between I1 and all other phalanges. Note that all subjects expect Subject 1 show an early response for at least one of the on-scalp magnetometers for I1. Compare with Fig. 5 in the manuscript.*

### Minimum

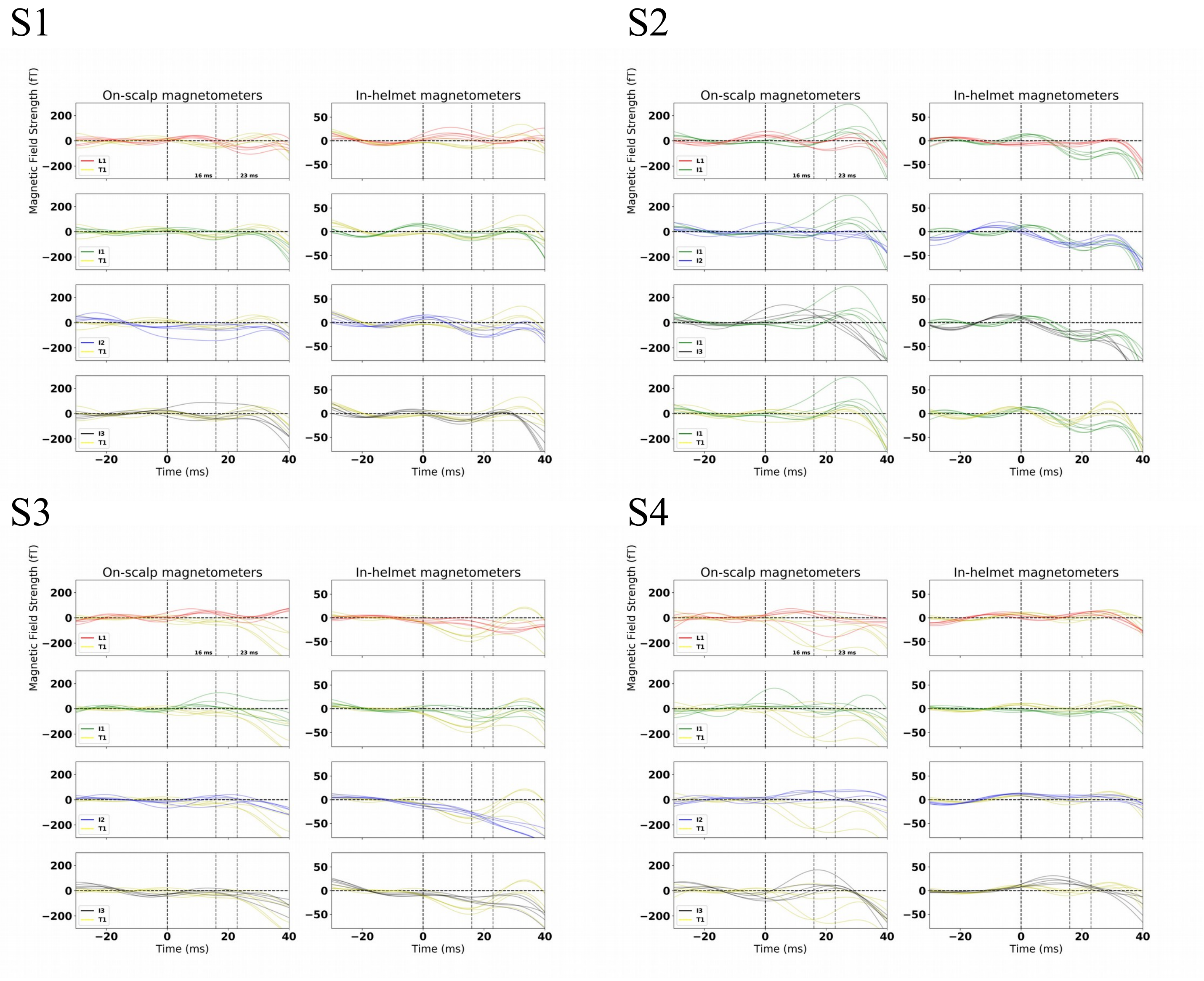


*Supplementary Fig. 12: Zoomed in view of the evoked responses (-30 to 40 ms) for the minimum position for the comparison between T1 and all other phalanges, except for Subject 2 for whom the comparison between I1 and all other phalanges is shown. Subjects 2 and 4 show early effects. Compare with Fig. 5 in the manuscript.*

**A**

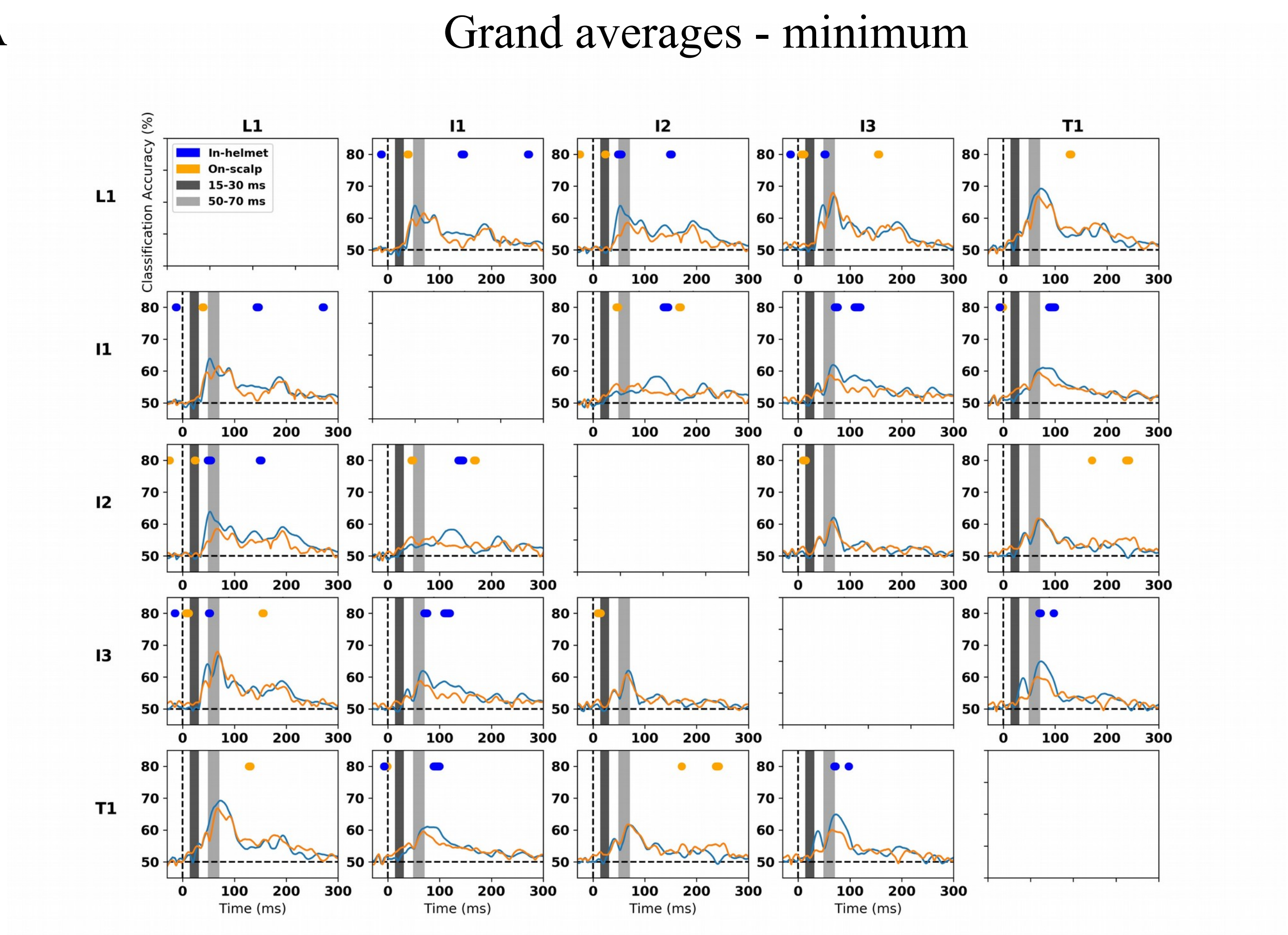


**B**

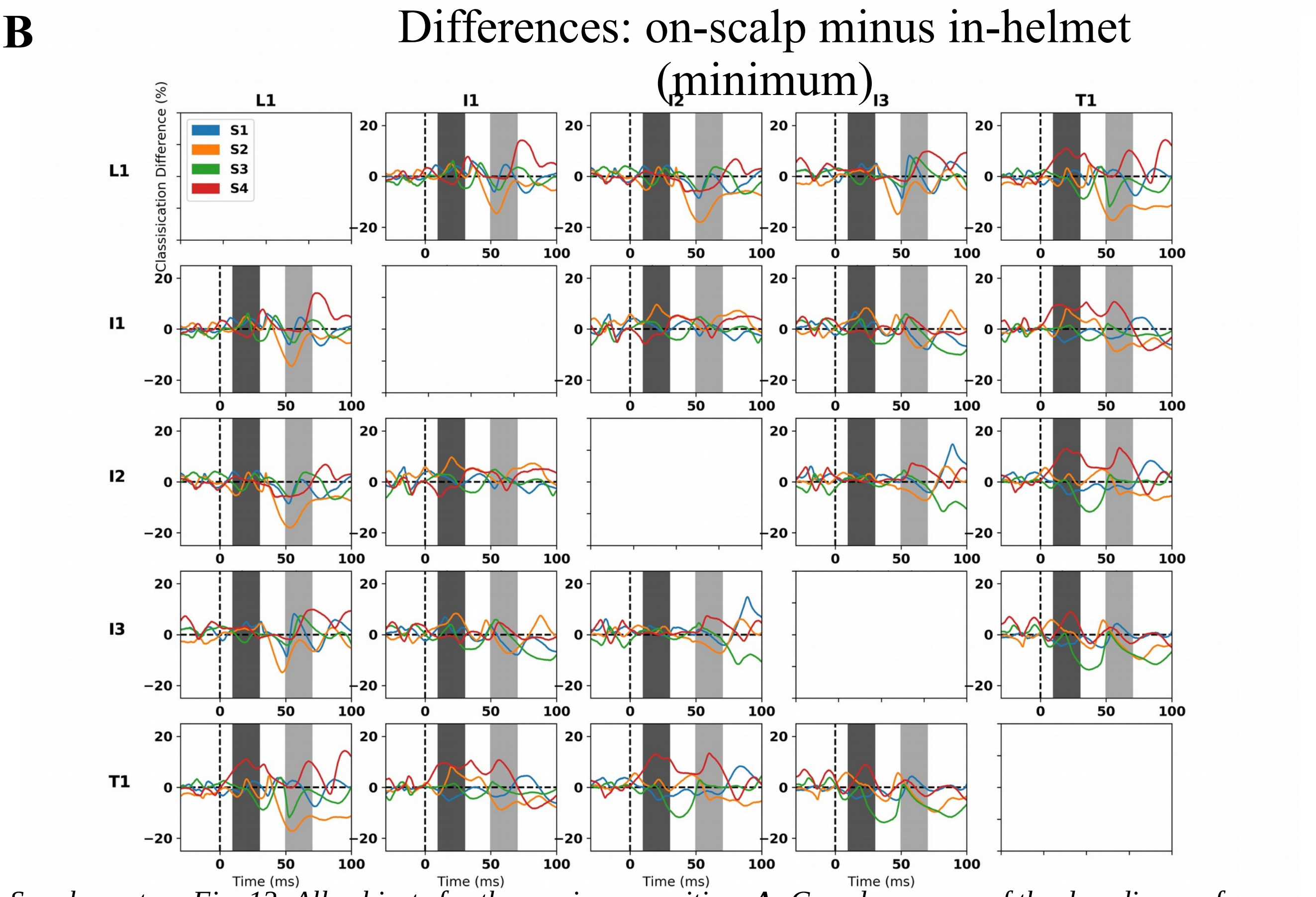


*Supplementary Fig. 13: All subjects for the maximum position.* ***A:*** *Grand averages of the decoding performance across all four subjects with dots indicating where classifications differed significantly (t-distribution with df=3) between on-scalp and in-helmet magnetometers. Orange dots indicate that on-scalp was better and blue dots that in-helmet was better. For the P60m component, it is seen that in-helmet measurements in general (significantly) outperforms on-scalp magnetometers.* ***B:*** *Differences between on-scalp and in-helmet classification. Subjects 2 and 4 show the greatest differences for the early time interval (15-30 ms) with on-scalp classifying the best.*

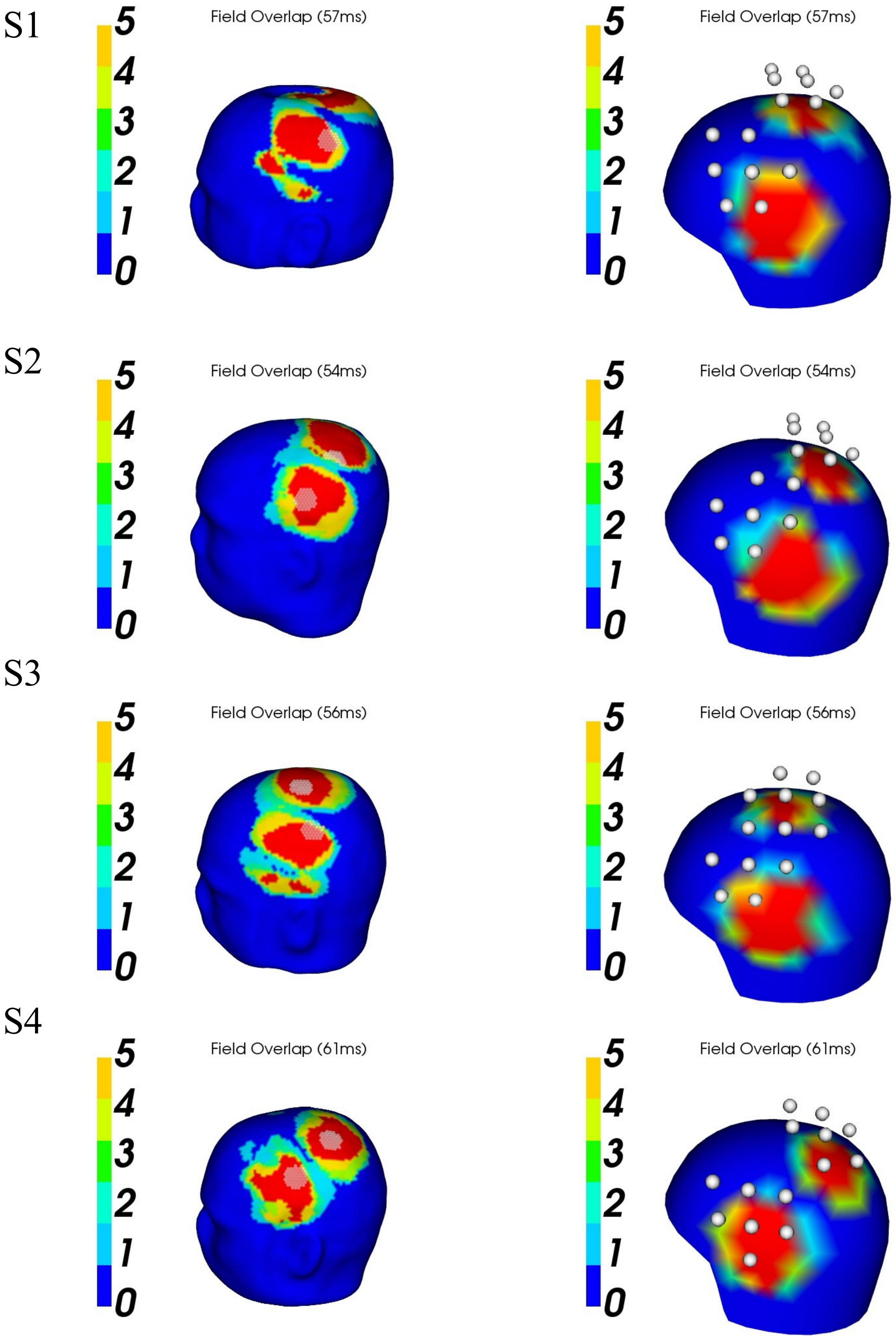


*Supplementary Fig. 14: The overlap of fields generated by the dipole models for each phalange at the mean peaking time over the five phalanges is illustrated. The lower and upper fifth percentiles of magnetic field strength on the head were calculated for* each *of these dipole models. The colour map indicates where these percentiles overlap. The white dots indicate the surface of the cryostat.*
